# A combined BSA-Seq and linkage mapping approach identifies genomic regions associated with Phytophthora root and crown rot resistance in squash

**DOI:** 10.1101/2020.09.13.295527

**Authors:** Gregory Vogel, Kyle E. LaPlant, Michael Mazourek, Michael A. Gore, Christine D. Smart

**Affiliations:** Plant Breeding and Genetics Section, School of Integrative Plant Science, Cornell University, Ithaca, NY 14853; Plant Pathology and Plant-Microbe Biology Section, School of Integrative Plant Science, Cornell University, Geneva, NY 14456

## Abstract

Phytophthora root and crown rot, caused by the soilborne oomycete pathogen *Phytophthora capsici*, leads to severe yield losses in squash (*Cucurbita pepo*). To identify quantitative trait loci (QTL) involved in resistance to this disease, we crossed a partially resistant squash breeding line with a susceptible zucchini cultivar and evaluated over 13,000 F_2_ seedlings in a greenhouse screen. Bulked segregant analysis with whole genome resequencing (BSA-Seq) resulted in the identification of five genomic regions – on chromosomes 4, 5, 8, 12, and 16 – featuring significant allele frequency differentiation between susceptible and resistant bulks in each of two independent replicates. In addition, we conducted linkage mapping using a population of 176 F_3_ families derived from individually genotyped F_2_ individuals. Variation in disease severity among these families was best explained by a four-QTL model, comprising the same loci identified via BSA-Seq on chromosomes 4, 5, and 8 as well as an additional locus on chromosome 19, for a combined total of six QTL identified between both methods. Loci, whether those identified by BSA-Seq or linkage mapping, were of small to moderate effect, collectively accounting for 28-35% and individually for 2-10% of the phenotypic variance explained. However, a multiple linear regression model using one marker in each BSA-Seq QTL could predict F_2:3_ disease severity with only a slight drop in cross-validation accuracy compared to genomic prediction models using genome-wide markers. These results suggest that marker-assisted selection could be a suitable approach for improving Phytophthora crown and root rot resistance in squash.

## Introduction

Squash and pumpkin (*Cucurbita pepo, C. maxima,* and *C. moschata*) are important crops grown for fruit that can be consumed either immature (e.g. summer squash) or mature (e.g. winter squash and pumpkin), as well as used decoratively (e.g. pumpkin and gourds). One of the major diseases affecting their production in the United States, which was valued at over $380 million in 2018 (United States Department of Agriculture 2019), is Phytophthora root and crown rot, caused by the soilborne oomycete *Phytophthora capsici*. Phytophthora root and crown rot causes lesions on roots and lower stems, leading to damping off in young seedlings and reduced vigor or death in older plants (Hausbeck and Lamour 2004). Disease management strategies are mostly limited to cultural practices meant to improve drainage, prevent splashing of infested soil, and reduce water movement within a field, in addition to the use of fungicides (Granke et al. 2012). However, the efficacy of several chemical active ingredients has been threatened by the emergence of fungicide insensitivity in pathogen populations (Lamour and Hausbeck 2000; Parra and Ristaino 2001; Dunn et al. 2010; Jackson et al. 2012). Host resistance is greatly desired by growers but has not been identified in appreciable levels in commercial cultivars.

To date, no sources of complete Phytophthora root and crown rot resistance have been described in squash, although germplasm screens in *C. pepo* (Padley et al. 2008) and *C. moschata* (Chavez et al. 2011) revealed a range of resistance levels among accessions of both species. In *C. pepo*, the most economically valuable of the three major domesticated squash species (Lust and Paris 2016), cultivars of *C. pepo* ssp. *pepo* (e.g. zucchini and pumpkin) are slightly less susceptible than cultivars of *C. pepo* ssp. *ovifera* (e.g. crookneck summer squash and acorn squash), but these differences are too small to provide economically relevant levels of disease control (Camp et al. 2009; Meyer and Hausbeck 2012; Krasnow et al. 2017). Little is known about Phytophthora root and crown rot resistance in wild *Cucurbita* species, which have played an important role in breeding for resistance to other diseases of squash (Rhodes 1964; Menezes et al. 2015; Holdsworth et al. 2016), although a *C. moschata* breeding line with introgressions from *C. lundelliana* and *C. okeechobeenesis* subsp. *okeechobeenesis* showed resistance to several Florida isolates of *P. capsici* (Padley et al. 2009). In crosses with different breeding lines representing distinct sources of resistance, segregation ratios suggest that resistance is genetically complex and controlled by multiple genes (Padley et al. 2009; Michael et al. 2019).

Selection for Phytophthora root and crown rot resistance is typically performed in controlled environments at the seedling stage (Padley et al. 2009; LaPlant et al. 2020). Nevertheless, predictable levels of disease pressure are often difficult to obtain, as the extent and rate of symptom development are highly influenced by the choice of pathogen isolate and inoculation method as well as the environmental conditions in the greenhouse and age of plants at the time of inoculation (Lee et al. 2001; Tian and Babadoost 2004; Enzenbacher and Hausbeck 2012). Higher than expected disease severity in a selection screen may leave little genetic variation remaining among the survivors in a breeding population and result in the death of plants possessing desirable resistance alleles. Knowledge of the quantitative trait loci (QTL) conferring Phytophthora root and crown rot resistance would enable marker-assisted selection (MAS), eliminating or reducing the number of generations in which plants would need to be inoculated in a resistance breeding program. While QTL conferring Phytophthora root and crown rot resistance have been reported recently in *C. moschata* (Ramos et al. 2020), to our knowledge none have been identified in *C. pepo*.

The combination of bulked segregant analysis with whole genome resequencing (BSA-Seq), first shown in yeast to be capable of detecting QTL of both major and minor effect (Ehrenreich et al. 2010; Wenger et al. 2010), has since become a popular method for mapping QTL in crop species (Takagi et al. 2013; Yang et al. 2013; Lu et al. 2014; Illa-Berenguer et al. 2015). Compared to traditional linkage mapping, where progeny are individually genotyped and phenotyped, BSA requires the preparation of fewer samples for genotyping and allows for more efficient phenotyping, as only individuals representing the phenotypic extremes of a population need to be identified (Michelmore et al. 1991). The choice of whole genome resequencing as the genotyping strategy in BSA provides an effective way to estimate genome-wide allele frequencies in bulks (via allele-specific read counts) without the need for prior marker development (Ehrenreich et al. 2010; Magwene et al. 2011; Takagi et al. 2013). BSA-Seq, however, features several disadvantages compared to traditional QTL mapping, such as the inability to estimate allelic effect sizes or to test for QTL by QTL interactions (i.e. epistasis).

In this study, we used a combination of BSA-Seq and traditional linkage mapping to discover QTL conferring resistance to Phytophthora root and crown rot in a biparental zucchini (*C. pepo*) population. Using a cross between a susceptible zucchini cultivar and a Cornell breeding line with intermediate resistance to Phytophthora root and crown rot, but poor fit as a zucchini or summer squash cultivar due to its growth habit and fruit shape (LaPlant et al. 2020), our goal was to identify QTL that can be directly used in MAS for introgressing disease resistance loci into genetic backgrounds representing more widely grown squash types. By synthesizing results from the two QTL mapping methods, we were able to estimate allelic effect sizes, generate lists of candidate genes in QTL regions, and test the predictive ability of QTL in independent datasets from those used for their detection.

## Materials and Methods

### Development of mapping population

A replicated greenhouse experiment evaluating germplasm from the Cornell squash breeding program for Phytophthora root and crown rot resistance resulted in the identification of one F_4:5_ family, Pc-NY21, with superior resistance compared to the other entries in the trial (LaPlant et al. 2020). This family was selected from a cross between Romulus, a Cornell zucchini cultivar, and PI 615089 from the National Plant Germplasm System, a white vegetable marrow accession with partial Phytophthora resistance. Eight individual F_5_ progeny from Pc-NY21, which were descended from the same F_4_ plant, were crossed with ‘Dunja F_1_’ (Enza Zaden), a susceptible zucchini variety, and an equal number of individuals from the eight F_1_ families were then intermated to generate an F_2_ mapping population.

### Selection of bulks and sequencing of DNA pools

Two separate sets of F_2_ individuals (Rep 1 and Rep 2) were evaluated for root and crown rot resistance in a greenhouse at Cornell AgriTech in Geneva, NY, in the fall of 2017. For each set, 6,912 seeds were sown in 72-cell trays (for a total of 96 trays) and inoculated with a zoospore suspension of *P. capsici* when the plants had two fully expanded true leaves (17-18 days after sowing). New York *P. capsici* isolate 0664-1 (Dunn et al. 2010) was used for inoculations and inoculum was prepared as in LaPlant et al. (2020). Inoculum was diluted to a concentration of 1×10^4^ zoospores/mL and sprayed over the tops of trays using a diaphragm pump backpack sprayer at a rate so as to deliver an intended total of 144 mL per tray, for a target of 20,000 zoospores (2 mL) inoculated onto each seedling.

A random (RAN), susceptible (SUS), and resistant (RES) tissue bulk were selected in each replicate. In order to target 15% of the population to include in each bulk, as has been shown in simulations to maximize QTL detection power and resolution (Magwene et al. 2011; Takagi et al. 2013), approximately 11 plants were sampled from each 72-cell tray. Plants for the RAN bulks were randomly sampled within 3 days prior to inoculation. Plants for the SUS bulks, which were sampled 4-5 days after inoculation, were visually identified as those within their respective trays featuring the greatest degree of wilt, stem necrosis, and sporulation on stem lesions. Plants for the RES bulks were sampled 7-10 days after inoculation and were identified as the plants that featured the least degree of wilt and leaf chlorosis in their respective trays. In trays where the most phenotypically extreme 11 plants were difficult to distinguish, as few as 6 or as many as 16 plants were sampled for the SUS and RES bulks. Tissue was sampled from all selected plants by taking a 3 mm hole punch from a newly emerging leaf. Tubes containing tissue samples were flash frozen in liquid nitrogen, stored at -80° C, and then lyophilized for 48 h prior to DNA extraction.

To minimize the heterogeneity of the DNA contributed by each individual to its pool, tissue samples from each tray were bulked separately for a total of 96 tubes per bulk (RAN, SUS, or RES). After extracting DNA from samples representing within-tray bulks, equimolar volumes of DNA from the 96 samples were then combined for a final DNA pool representing selections from all trays. This strategy was elected as a compromise between performing a separate DNA extraction for every individual plant sample and bulking >1,000 leaf punches prior to DNA extraction. DNA extractions were performed using the DNeasy 96 plant kit (Qiagen, Valencia, CA, USA), with the following modifications to maximize the purity and integrity of DNA from diseased plant samples: the volumes of buffers AP1, P3, and AW1 were doubled; the -20° C incubation was extended from 20 to 60 m; and an additional wash step was performed with 800 μl molecular-grade ethanol. The DNA concentration of each sample was measured using a Qubit fluorometer (Thermo Fisher Scientific, Waltham, MA, USA) prior to pooling.

The six DNA pools (RAN, SUS, and RES for each of the two replicates) and parental DNA samples were submitted to the Cornell University Biotechnology Resource Center for library preparation and sequencing. To represent the resistant parent, DNA was extracted from the F_4_ progenitor of Pc-NY21. PCR-free libraries were created by mechanically shearing DNA samples and ligating adapters, using reagents equivalent to those in the TruSeq library preparation kit (Illumina, San Diego, California). For several DNA pools, two separate libraries were created, resulting in technical replicates. Library yields were determined using digital droplet PCR. Libraries were then pooled and sequenced on either 1 or 2 lanes, depending on the library, of an Illumina NextSeq500 generating paired-end 150 bp reads. Due to a technical error during library preparation, reads for Rep 1 RAN were unusable and therefore not included in any analyses.

### BSA-Seq data analysis

Raw reads were filtered and trimmed from both ends using fastp v 0.20.0 (Chen et al. 2018) with default parameters, except the arguments --*correction* and -- *trim_poly_g* were enabled to correct bases in overlapped regions and remove polyG strings in read tails. Trimmed reads were then aligned to the *C. pepo* reference genome (v 4.1; Montero-Pau et al. 2018) with bwa v 0.7.17 (Li and Durbin 2009) using the MEM algorithm and default parameters. Resulting bam files were sorted and indexed with samtools and variants were called using the bcftools *mpileup* and *call* commands (Li et al. 2009). The minimum alignment mapping quality was set to 20 and the consensus-caller method was used. The resulting VCF file was filtered with VCFtools version 0.1.17 (Danecek et al. 2011) to remove indels and SNPs with more than 2 alleles. Reference and alternate allele counts for every SNP in each pool were then extracted from the filtered VCF file using a custom python script, and imported into R (R Core Team 2019) for filtering.

SNPs with a total read depth lower than the 5^th^ percentile or higher than the 95^th^ percentile of the read depth distribution across all SNPs were removed, as were SNPs with a minor allele frequency (calculated using allele counts in each pool) <0.10. Unanchored SNPs (i.e. those on scaffold Cp4.1LG00) were also removed. Parental genotypes with fewer than 6 reads were set to missing, and only SNPs where the parents were homozygous for opposite alleles were retained. Parental genotypes were then used to recode alleles from reference/alternate coding to Dunja-derived/Pc-NY21 derived. Allele counts in samples representing technical sequencing replicates of the same DNA pool were summed at this point. Further filtering was then performed within each pool, by setting to missing any sites with an allele frequency <0.10 or >0.90 or with a read depth lower than the 10^th^ percentile for that pool. These missing sites were not included for calculation of allele frequencies nor for any statistical test involving that pool.

For visualization of allele frequencies, allele frequency means for each pool were estimated in 500 Kb sliding windows with a 100 Kb increment. To test for deviations in allele frequencies between SUS and RES pools, the software program MULTIPOOL (v 0.10.2; Edwards and Gifford 2012), which estimates pool allele frequencies using a dynamic Bayesian network, was used. The argument –mode was set to *contrast*. We set 1,056 as the number of individuals contributing DNA to each pool (-n) and 125,636 as the length, in base pairs, of a centimorgan in squash (-c), which was estimated by dividing the mean of the lengths of three published genetic maps (Esteras et al. 2012; Holdsworth et al. 2016; Montero-Pau et al. 2017) by the estimated *C. pepo* genome size of 283 Mb (Montero-Pau et al. 2018). MULTIPOOL was also used to test for allele frequency deviations between Rep 1 SUS and Rep 2 SUS as well as Rep 1 RES and Rep 2 RES in order to generate a null distribution of LOD scores reflecting the comparison between two independent bulks of plants selected in the same direction.

### Phenotyping of F_2:3_ population

One hundred eighty-seven F_2_ plants were self-pollinated to generate F_2:3_ families. These F_2_ plants represented two different cohorts of individuals: a ‘random’ cohort of 169 plants started from remnant seed and a ‘selected’ cohort comprising 18 survivors from the BSA-Seq screen. In order to collect seed from infected plants from the BSA-Seq screen, 42 seedlings (18 in Rep 1 and 24 in Rep 2) that appeared healthy at 14-15 days post inoculation were treated with mefenoxam (Ridomil Gold EC; Syngenta AG, Basel, Switzerland) and transplanted to 3-gal pots. Eighteen of these 42 plants survived and produced viable seed after self-pollination.

The 187 F_2:3_ families, in addition to parental and F_1_ checks, were evaluated for Phytophthora root and crown rot resistance as seedlings in the greenhouse. An F_5:6_ family derived from one selfed F_5_ Pc-NY21 individual was included to represent the resistant parent. Experimental units consisted of 12 adjacent cells in a 72-cell tray and were arranged in a randomized complete block design with three replications. Three of the 187 families were only included in two blocks due to limited seed. Inoculum was prepared as in the BSA-Seq screen, except plants were inoculated by pipetting a suspension of 10,000 zoospores to the potting soil surface adjacent to each plant.

Plots were rated for incidence of mortality at several days post inoculation (dpi) (3, 5, 7, 10, and 12 dpi for Rep 1; 3, 4, 5, 6, 7, and 10 dpi for Reps 2 and 3). Seedlings were declared dead when they either had all wilted leaves, had only one or fewer non-chlorotic leaves, or were completely prostrate due to stem lesions. The relative Area Under the Disease Progress Curve (rAUDPC; Fry 1978) was then calculated for each plot according to the following formula:

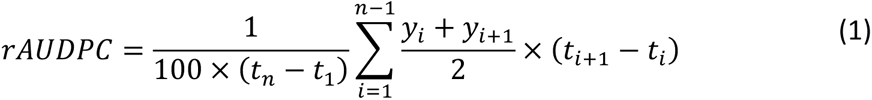

where *y*_i_ is the mortality rating as a percentage at the *i*th observation, *t*_i_ is the time point in days at the *i*th observation, and *n* is the total number of observations. The relative AUDPC was used to normalize AUDPC values between blocks as they were rated for different numbers of days.

Ratings for any plot with fewer than 6 germinated plants were set to missing. The following mixed linear model was then fit using the R package lme4 (Bates et al. 2007) in order to estimate best linear unbiased estimators (BLUEs) for the effect of each family on rAUDPC:

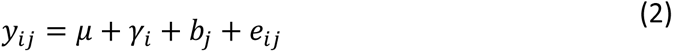

In this model, *y_ij_* are raw phenotypic observations, *μ* is the grand mean, *γ_i_* is the fixed effect for the *i*th family, 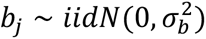 is a random effect for the *j*th block (i.e. replication), and 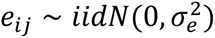 is the residual effect. Separate models were also fit using mortality ratings at 3, 5, 7, and 10 dpi as the response variables. In order to calculate line-mean heritability (*H*^2^; Holland et al. 2003), a similar model was fit, except family was included as a random instead of fixed effect. The following formula was then used:

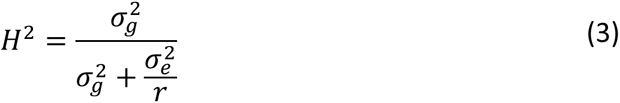

where 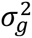 is the genotype variance, 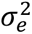 is the error variance, and *r* is the harmonic mean of the number of replications per family without missing data.

### Genotyping of F2 individuals and genetic map construction

Newly emerged leaves were sampled from the F_2_ parents of the F_2:3_ mapping population and desiccated using silica gel (Millipore Sigma, Burlington, MA). Total genomic DNA was then isolated using the DNeasy Plant Mini Kit (Qiagen, Valencia, CA, USA) following manufacturer’s directions, except DNA was eluted into ultrapure water instead of Buffer AE. DNA samples were obtained from a total of 188 F_2_ progeny (19 ‘selected’ and 169 ‘random), as well as the two parents of the population, with the F_4_ progenitor of Pc-NY21 sampled to represent the resistant parent. DNA samples were sent to the University of Wisconsin-Madison Biotechnology Center for preparation of genotyping-by-sequencing (GBS) libraries digested with *Ape*KI (Elshire et al. 2011). Libraries were then paired-end sequenced on an Illumina NovaSeq6000 with a target output of two million reads per sample.

Genotypes were called using GBS-SNP-CROP (Melo et al. 2016), a variant calling pipeline designed to run with paired-end reads. As part of this pipeline, reads were demultiplexed, trimmed using Trimmomatic (v 0.39; Bolger et al. 2014), and aligned to the squash reference genome (Montero-Pau et al. 2018) using BWA-mem (Li and Durbin 2009). Variants were called using the default parameters in GBS-SNP-CROP, except the arguments -altStrength, - mnAlleleRatio, -mnAvgDepth, and -mxAvgDepth were set to 0.9, 0.15, 7, and 150, respectively. Genotype calls were then converted to a dosage matrix and imported into R for filtering.

Unanchored SNPs were removed, as were SNPs that did not meet the following criteria: minor allele frequency (MAF) >0.1, call rate >0.9, and read depth between 15 and 90. Three individuals with over 40% missing data were removed as well. SNPs were then removed that were not homozygous for opposite alleles in the parents or that failed a chi-squared goodness of fit test (p < 0.01) for 1:2:1 AA:AB:BB segregation among the F_2_ progeny. In order to identify potential errors during DNA extraction or library preparation, pairwise genotype concordance was calculated among all pairs of samples and four pairs of samples were found to have >99% identical genotype calls. For each pair, the sample with the least missing data was retained for genetic map construction and the phenotypic records for those retained samples were set to missing. Finally, markers were pruned by identifying sequences of consecutive SNPs with identical genotype calls and retaining the marker with the least missing data. Genotypes were then re-coded as “A/B/H” for genetic map construction.

The R package R/qtl (Broman et al. 2003) was used to estimate recombination fractions between all pairs of markers in order to build a genetic map. Linkage group assignments were based on the physical coordinates of markers on the reference genome, except for six markers which displayed high recombination fractions with all markers on their respective linkage groups and were re-located accordingly to the correct linkage group. The MSTMap algorithm (Wu et al. 2008) was then used to re-order markers within each linkage group, using the R package ASMap (Taylor and Butler 2017). Recombination fractions were converted into map distances using the Kosambi map function.

### Linkage mapping and validation of BSA-Seq QTL

A multiple QTL mapping (MQM) procedure was performed in R/qtl (Broman and Sen 2009) using the 176 F_2:3_ families with genotype and phenotype data that passed filtering. First, conditional genotype probabilities were calculated at 1 cM positions across the genetic map using the *calc.genoprob* function. The genotyping error rate was set to 0.001, reflecting the average proportion of discordant genotype calls among the four pairs of erroneously duplicated samples. In order to estimate penalties for the inclusion of terms in model selection, rAUDPC values were permuted 1,000 times relative to genotypes and a two-dimensional QTL scan using Haley-Knott regression (Haley and Knott 1992) was performed using the *scantwo* function. Penalties at *α*=.05 were derived from the resulting null distribution of LOD scores using the *calc.penalties* function, and the best fitting QTL model was identified with the forward/backward stepwise search algorithm implemented in the *stepwiseQTL* function, again using Haley-Knott regression. The function *bayesint* was used to calculate 90% Bayesian credible intervals for QTL locations, with the ends of the intervals extended to marker positions so as to convert genetic markers to physical coordinates on the reference genome.

The *fitqtl* function, which performs a likelihood ratio test after dropping one QTL from the model at a time, was used to calculate LOD scores and p-values for each QTL, as well as estimate their effect sizes and proportion of phenotypic variance explained. Two QTL models were fit, one containing the loci detected via BSA-Seq and one containing those identified from multiple QTL mapping using the F_2:3_ phenotypes. The QTL detected via BSA-Seq were placed on the genetic map by identifying, for each QTL, the GBS marker nearest the midpoint of the MULTIPOOL LOD peaks from Reps 1 and 2. For plotting QTL effects and estimating QTL allele frequencies in the two F_2_ cohorts, missing GBS genotypes were imputed using the Viterbi algorithm as implemented in the *argmax.geno* function. Significant differences between rAUDPC means for F_2:3_ families with differing QTL marker alleles were determined using Tukey’s honestly significant difference test at *α*=.05 using the function *HSD.test* in the R package agricolae (Mendiburu and Simon 2015).

### Synteny with *C. moschata*

To assess if QTL in *C. pepo* were syntenic with those reported by Ramos et al. (2020) in *C. moschata*, we followed the approach used in the ‘SyntenyViewer’ module of the Cucurbit Genomics Database (Zheng et al. 2019) in order to identify and visualize regions of synteny between the genomes of the two species. Briefly, the *C. pepo* (Montero-Pau et al. 2018) and *C. moschata* (Sun et al. 2017) protein sequences were aligned against each other using blastp (Camacho et al. 2009) and synteny blocks were identified from the resulting alignments using MCScanX (Wang et al. 2012). The R package circlize (Gu et al. 2014) was then used to visualize syntenic relationships.

### Candidate gene identification

QTL regions, which we defined as the union of BSA-Seq and MQM credible intervals, were cross-referenced with the squash reference gff file (Montero-Pau et al. 2018) using bedtools v 2.28 (Quinlan and Hall 2010) in order to generate a list of genes annotated in QTL regions. SnpEff v 4.3T Cingolani et al. 2012) was then used to annotate variants between Pc-NY21 and Dunja for their putative functional impact on these genes. The whole genome resequencing genotype calls from BSA-Seq were used for variant annotation. These genotype calls were filtered on read depth and minor allele count frequency as for the BSA-Seq analysis, except, in order to prevent the removal of potential causative variants, indels were retained, low read-depth sites were not set to missing, and variants that were heterozygous in one parent were not removed. Variants where the minor allele was not called in at least 5 samples, including parents and pools, were removed. Genes in QTL regions were also evaluated for their homology with melon (*Cucumis melo*) genes shown to be differentially expressed following inoculation with *P. capsici*, as reported by Wang et al. (2020). Homologues were defined as reciprocal best hits on the basis of e-value after using blastp (Camacho et al. 2009) to align the complete sets of melon (v 3.6.1; Garcia-Mas et al. 2012) and squash (Montero-Pau et al. 2018) protein sequences against each other.

### Prediction models

A cross-validation approach was used to assess the ability of different linear models, featuring markers either genome-wide or only in QTL regions, to predict rAUDPC estimates among F_2:3_ families. Four models were compared: a multiple regression model in which the allele dosages at the GBS markers tagging each BSA-Seq QTL were treated as fixed effects (QTL MLR); a genomic best linear unbiased prediction model (GBLUP), where genome-wide markers were used to model the covariance between families with a genomic relationship matrix; a GBLUP plus QTL model (GBLUP+QTL) where, in addition to the genomic relationship matrix, QTL marker allele dosages were included as fixed effects; and a whole-genome regression Bayesian approach (BayesB; Meuwissen et al. 2001), in which marker effects are assigned a prior distribution where a portion equal zero and the rest have a scaled-*t* distribution. The QTL MLR model was fit using the lme4 R package (Bates et al. 2007), the two GBLUP models using R package rrBLUP (Endelman 2011), and the BayesB model using R package BGLR (Perez and Gustavo de los Campos 2014). The genotype data used in models, whether genome-wide or only at the five QTL markers, were from the genetic map-imputed GBS SNP set.

For cross-validation, the data were partitioned randomly into two subsets: a training set used to fit a given model, comprising 80% of the individuals, and a test set used for assessing prediction accuracy, comprising 20% of the individuals. The same partitions were used for all five models, to enable direct comparison, and this was replicated 50 times. In each cross-validation partition, prediction accuracy was estimated by taking the Pearson’s correlation coefficient between the observed values and estimated breeding values of the test individuals, as predicted by the training-set model.

### Data availability

Whole genome resequencing reads used in BSA-Seq and genotyping-by-sequencing reads used in linkage mapping have been deposited in the National Center of Biotechnology Information Sequence Read Archive (SRA) under BioProject accession number PRJNA662576. Files containing allele counts used in BSA-Seq, genotype and phenotype data used in linkage mapping, and annotated variant calls used for candidate gene identification are available at CyVerse (https://datacommons.cyverse.org/browse/iplant/home/shared/GoreLab/dataFromPubs/Vogel_SquashQTL_2020). All scripts used for data analysis are available on Github (http://github.com/gmv23/Pcap-QTL-Mapping).

## Results

### BSA-Seq

DNA pools were sequenced from two independent replicates of resistant, susceptible, and random (Rep 2 only) bulks selected from phenotypic screens featuring >6,500 F_2_ squash seedlings each. A total of 192 Gb of sequencing reads were generated from the five DNA pools and the parents of the population (Table 1). After aligning reads to the squash reference genome and identifying variants, between 182,311-186,020 SNPs were retained in each pool after filtering. These SNPs featured a median read depth of 9-10 in the parents, 57-70 in SUS and RES pools, and 24 in the Rep 2 RAN pool (Table 1). While the majority of the genome featured dense marker coverage, several regions contained few or no SNPs, including regions >1 Mb in size on chromosomes 7, 13, 14, 16, and 20 (Figure S1; Figure S2).

**Table 1.**
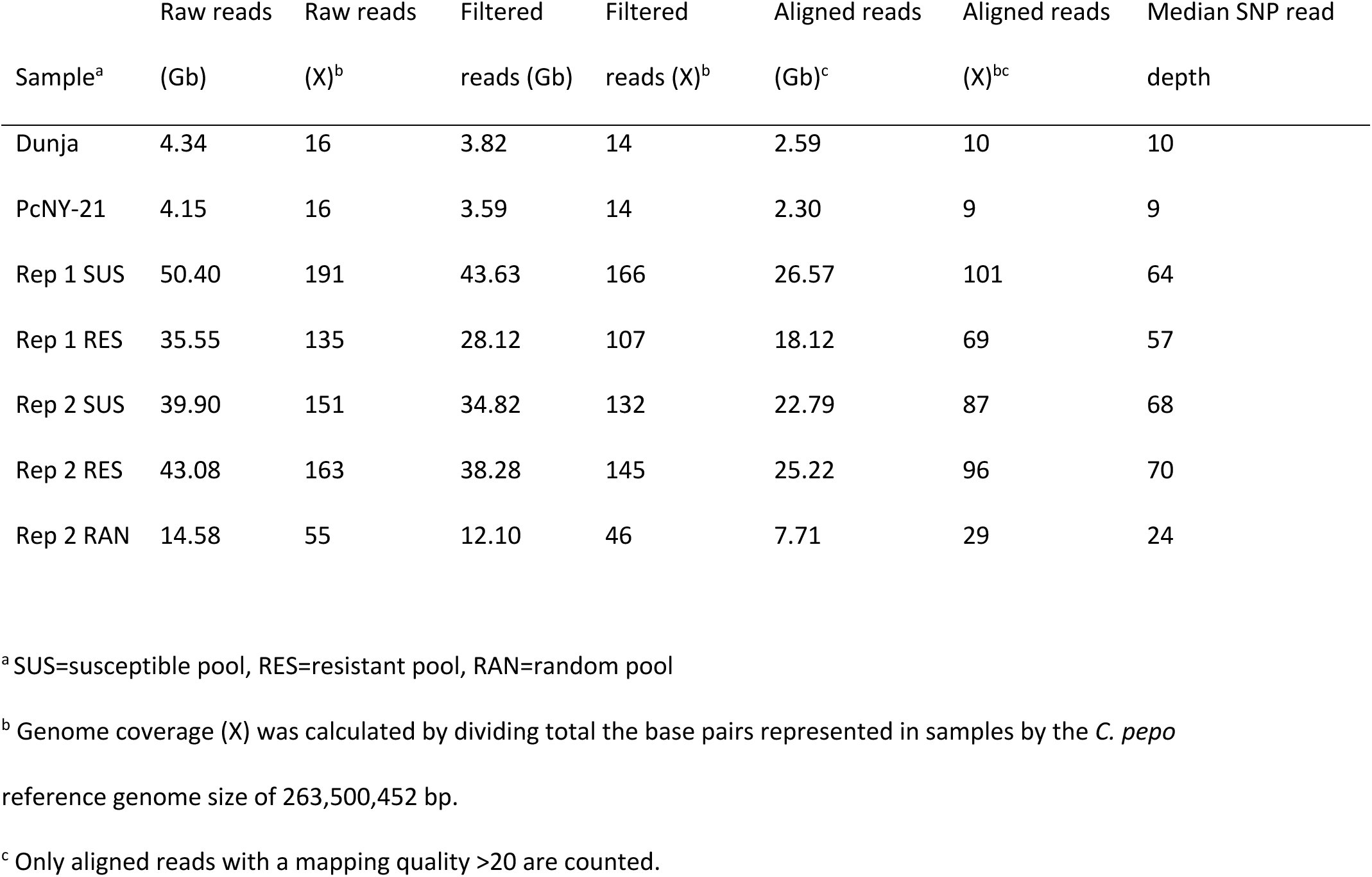
Sequencing statistics for BSA-Seq samples.

Allele frequencies at individual SNPs fluctuated, with the median standard deviation of SNP allele frequencies binned in 500 Kb windows ranging from 0.07 in Rep 2 RES, the pool with the highest read depth, to 0.11 in Rep 2 RAN, the pool with the lowest read depth. Averaging individual values in 500 Kb sliding windows enabled easier visualization of allele frequencies, showing that for the majority of the genome, RES, SUS, and RAN pools featured little differentiation (Figure S1; Figure S2).

We used the program MULTIPOOL to test for differences in allele frequencies between pools and determine 90% credible intervals for QTL positions. A null test between replicates for both RES and SUS pools resulted in a maximum LOD score of 2.59, and we decided to use a conservative LOD score threshold of 4 for QTL detection. Five genomic regions, on chromosomes 4, 5, 8, 12, and 16, featured LOD scores >4 in both Reps 1 and 2 after testing for differences between RES and SUS pools (Table 2; Figure 1). Two additional regions, on chromosome 17 and the beginning of chromosome 4, surpassed the LOD score threshold in only one of two reps, and were therefore not considered for further analysis. In the QTL regions on chromosomes 4, 5, and 8, the RES pool featured a higher frequency of the Pc-NY21 allele compared to the SUS pool, whereas the opposite was true in the chromosome 12 and 16 regions, indicating that the allele conferring resistance in each of these two regions was inherited from susceptible parent Dunja. In all five QTL regions, the RAN pool in Rep 2 displayed allele frequency values intermediate between those of the RES and SUS pools. Segregation distortion was evident in the chromosome 8 QTL, where RES, SUS, and RAN pools all showed enrichment for the Pc-NY21 allele, although this deviation was greater in the RES compared to the SUS pool.

**Figure 1.**
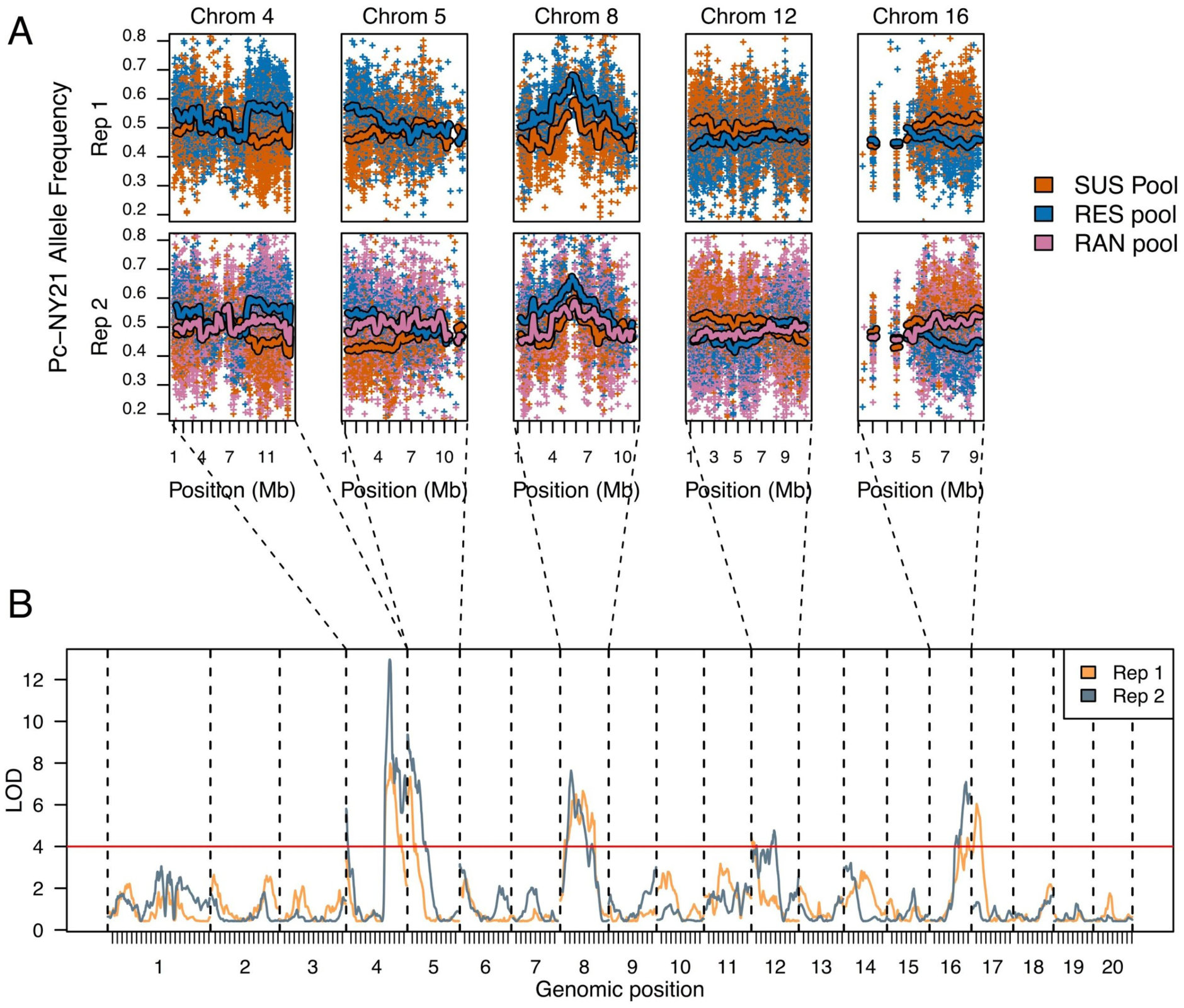
A) Pc-NY21 allele frequencies in susceptible (SUS), resistant (RES), and random (RAN) (Rep2 only) pools for all chromosomes that feature a region surpassing a MULTIPOOL LOD score of 4 in both reps. Plus signs are individual SNP allele frequencies and lines are smoothed means calculated in 500 Kb sliding windows with a 100 Kb increment. For ease of visualization, only a random subset of 25% of SNPs are shown. Smoothed means were not calculated in any window featuring fewer than 30 SNPs. B) Genome-wide LOD scores from MULTIPOOL testing for allele frequency deviations between susceptible and resistant pools for Reps 1 and 2 of BSA-Seq.

**Table 2.** BSA-Seq QTL mapping results.

| Chromosome | Max LOD<sup>ab</sup> | Max LOD Position<br>(Mb) (90% CI)<sup>ab</sup> | Max allele<br>frequency<br>difference<sup>bc</sup> | Max allele<br>frequency<br>difference position<br>(Mb)<sup>bc</sup> |
| --- | --- | --- | --- | --- |
|  |  | 9.16 (8.3-10.38) / |  |  |
| Cp4.1LG04 | 7.98 / 12.96 | 9.04 (8.7-9.36) | 0.13 / 0.17 | 9.25 / 9.05 |
|  |  | 0.6 (0.18-1.68) / |  |  |
| Cp4.1LG05 | 7.34 / 9.36 | 0.05 (0.04-2.47) | 0.12 / 0.13 | 0.65 / 2.55 |
|  |  | 4.7 (1.95-6.72) / |  |  |
| Cp4.1LG08 | 6.66 / 7.64 | 2.25 (1.84-5.18) | 0.12 / 0.13 | 3.15 / 2.55 |
|  |  | 0.06 (0.06-6.9) / |  |  |
| Cp4.1LG12 | 4.4 / 4.77 | 4.71 (0.36-7.38) | -0.09 / -0.09 | 0.25 / 4.95 |
|  |  | 6.36 (4-8.22) / 7.57 |  |  |
| Cp4.1LG16 | 4.6 / 7.09 | (5.84-8.26) | -0.10 / -0.12 | 7.95 / 7.45 |

<sup>a</sup> Maximum LOD scores, positions, and 90% credible intervals (CIs) determined using MULTIPOOL.
<sup>b</sup> Values for Reps 1 and 2 are separated by a “/.”
<sup>c</sup> Maximum smoothed allele frequency difference, calculated as mean resistant pool allele frequency – mean susceptible pool allele frequency in 500 Kb sliding windows with a 100 Kb increment.

The maximum smoothed allele frequency difference between RES and SUS pools was modest in all cases, ranging from 0.09 on chromosome 12 (averaging across reps) to 0.15 on chromosome 4 (Table 2). For the QTL on chromosomes 4 and 5, which featured the highest LOD scores, 90% credible intervals ranged from 0.66 to 2.43 Mb in size, and the location of LOD peaks were consistent between reps, displaying a difference of 0.12 Mb on chromosome 4 and 0.55 Mb on chromosome 5. The QTL on chromosomes 8, 12, and 16, on the other hand, had larger credible intervals ranging from 2.41 Mb to 7.01 Mb in size, and showed little consistency in the location of QTL peaks between reps, varying in the case of the chromosome 12 QTL by as much as 4.65 Mb, almost half the length of the chromosome.

### Validation of BSA-SEQ results and linkage mapping with F_2:3_ families

One hundred eighty-seven F_2_ plants – comprising 18 survivors of the BSA-Seq screen (‘selected’ plants) and 169 plants grown from remnant seed (‘random’ plants) – were individually genotyped with genotyping-by-sequencing and selfed to generate an F_2:3_ mapping population. In a greenhouse experiment, the family-mean heritabilities among F_2:3_ families for mortality at 3,5,7, and 10 dpi ranged from 0.40-0.76, with heritability increasing with dpi (Table S1). A summary statistic incorporating ratings at all time points, rAUDPC, featured a higher heritability (0.77) compared to mortality at any individual time point, and was therefore used for all future analyses. The distribution of BLUEs for rAUDPC among F_2:3_ families appeared slightly non-normal, with a long left (i.e. resistant) tail (Figure 2A). Transgressive segregation was observed, with two families displaying rAUDPC estimates lower than Pc-NY21 and 33 families displaying rAUDPC estimates higher than Dunja. Lower rAUDPC estimates were observed among F_2:3_ families derived from ‘selected’ plants, which featured a median of 60.69, compared to F_2:3_ families from ‘random’ plants, which featured a median rAUDPC estimate of 74.18 (Table S1; Figure 2B).

**Figure 2.**
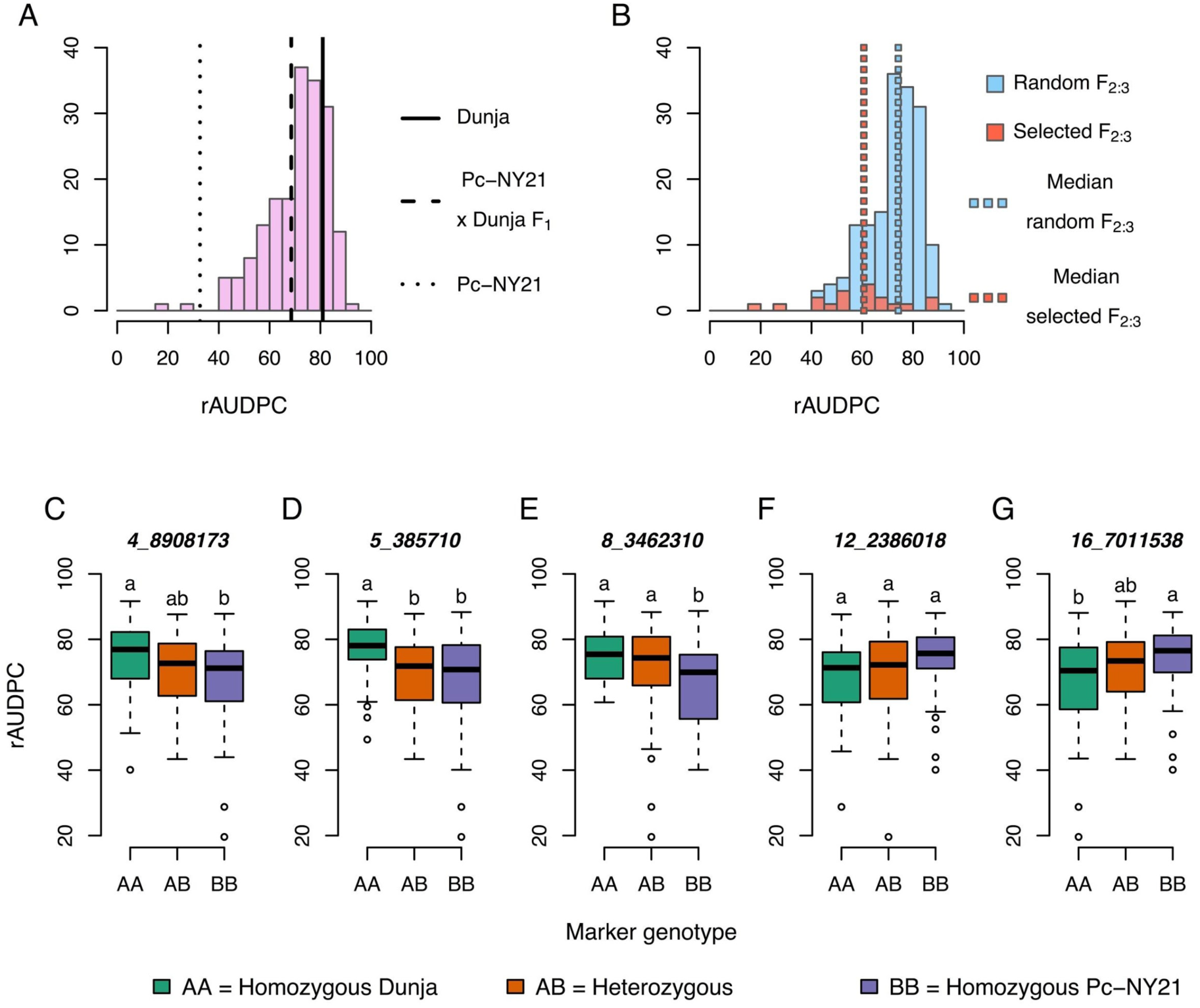
Distributions of best linear unbiased estimates of relative Area Under the Disease Progress Curve (rAUDPC) for 187 F_2:3_ families. A) Histogram of F_2:3_ rAUDPC estimates with lines representing estimates for the parents and F_1_ generation B) Overlapping histograms for F_2:3_ families derived from 169 ‘random’ F_2_ individuals grown from remnant seed (blue) and 18 ‘selected’ F_2_ individuals that survived the BSA-Seq screen. C-G) rAUDPC estimates for F_2:3_ families vs their F_2_ parent’s genotype at each of five SNP markers tagging QTL detected via BSA-Seq. Values are shown for 176 F_2:3_ families with both phenotype and genotype data.

A genetic map was constructed from 605 GBS-derived SNPs that were called on 181 F_2_ individuals with high-quality genotype data. The map was 2,023.38 cM in length and consisted of 20 linkage groups corresponding to the 20 chromosomes of *C. pepo* (Table S2; Figure S3). Linkage groups featured between 7-76 markers and ranged in length from 43.50-193.55 cM. Genetic marker positions were largely correlated with physical marker positions on the reference genome, except for 6 markers that mapped to the wrong linkage group and two genomic regions – covering approximately 8 Mb on the beginning of chromosome 4 and 1 Mb on the end of chromosome 17 – where the genetic marker order was inverted in relation to the physical marker order (Figure S4).

The GBS markers nearest the BSA-Seq QTL locations, which ranged from 1-196 Kb away from the midpoint of the LOD peaks from BSA-Seq Reps 1 and 2, were identified (Table S3). For all 5 of these QTL markers, the frequency of the resistant allele, as determined by BSA-Seq, was enriched in the ‘selected’ compared to the ‘random’ F_2:3_ families, with the magnitude of the allele frequency difference (0.19) greatest for the QTL on chromosome 4 and smallest (0.04) for the chromosome 16 QTL. rAUDPC means were significantly different among F_2:3_ families with different marker genotypes for the QTL on chromosomes 4, 5, 8, and 16, although not for the chromosome 12 QTL (Figure 3C). Consistent with the BSA-Seq results, the Pc-NY21 allele was associated with lower rAUDPC estimates at the chromosome 4, 5, and 8 QTL markers, and associated with higher rAUDPC estimates at the QTL markers on chromosomes 12 and 16. A multiple regression model fit with the 5 markers tagging BSA-Seq QTL explained 27.85% of the variation in rAUDPC estimates among F_2:3_ families, with individual QTL explaining between 1.94-9.84% of the phenotypic variation (Table 3). Additive allelic effects on rAUDPC ranged in absolute value from 1.93 for the chromosome 12 QTL to 4.98 for the QTL on chromosome 5, and dominance deviations, as a percentage of the additive effect, varied from 30-115%, indicating a mixture of partially dominant and dominant gene actions.

**Figure 3.**
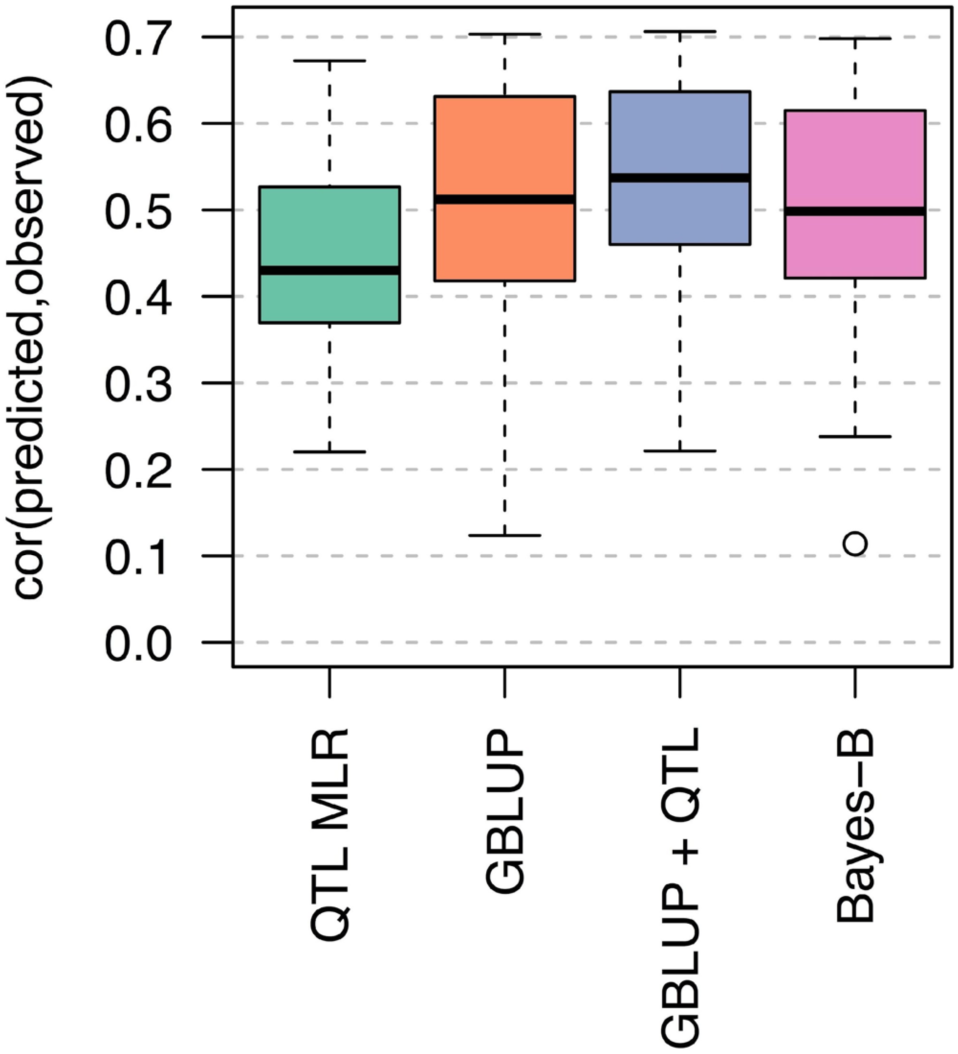
Cross-validation prediction accuracies for four models predicting F_2:3_ rAUDPC. QTL MLR: multiple linear regression model with allele dosages at five SNP markers tagging BSA-Seq QTL. GBLUP: Genomic best linear unbiased prediction model using 605 genome-wide SNPs. GBLUP+QTL: GBLUP model with five SNP markers tagging BSA-Seq QTL as fixed effects. Bayes-B: Bayes-B model using 605 genome-wide SNPs.

**Table 3.** Effect sizes of BSA-Seq QTL in F_2:3_ population.

| Chromosome | Position of QTL-tagging GBS | Position of QTL-tagging GBS | LOD | Proportion |  |  |  |
| --- | --- | --- | --- | --- | --- | --- | --- |
|  | marker (Mb)<sup>a</sup> | marker (cM)<sup>a</sup> |  | variance<br>explained<br>(%) | F test p-value | a<sup>b</sup> | d<sup>c</sup> |
| 4 | 8.91 | 84.16 | 2.98 | 5.85 | 0.00161 | -4.1 | 1.21 |
| 5 | 0.39 | 1.71 | 4.88 | 9.84 | 2.64E-05 | -4.98 | -3.4 |
| 8 | 3.46 | 44.48 | 3.11 | 6.11 | 0.00122 | -4.01 | 2 |
| 12 | 2.39 | 24.94 | 1.18 | 2.26 | 0.0787 | 1.93 | -2.22 |
| 16 | 7.01 | 49.84 | 1.01 | 1.94 | 0.112 | 2.06 | 1.63 |

<sup>a</sup> Genotyping-by-sequencing (GBS) markers representing BSA-Seq QTL were those nearest the midpoint of MULTIPOOL LOD peaks from BSA-Seq Reps 1 and 2.
<sup>b</sup> a=additive effect of Pc-NY21 allele on rAUDPC.
<sup>c</sup> d=dominance effect of Pc-NY21 allele.

The genetic map and F_2:3_ phenotypic data were also used to discover QTL *de novo* using a multiple QTL mapping approach. The best fitting model, explaining 35.17% of the phenotypic variation, was found to contain four non-interacting QTL, consisting of the same loci identified via BSA-Seq on chromosomes 4, 5, and 8, in addition to a locus on chromosome 19 where the resistant allele was contributed by Dunja (Table 4; Figure S5). MQM and BSA-Seq credible intervals overlapped for the three QTL identified via both approaches, except for the chromosome 4 QTL, where the BSA-Seq Rep 2 and MQM credible intervals were disjoint (Figure 4). For the QTL on chromosomes 5 and 8, the most likely QTL positions as determined by MQM were very close to the GBS markers tagging BSA-Seq QTL, differing by 0.48 cM on chromosome 8 and falling on the same marker on chromosome 5. In the case of the chromosome 4 QTL, however, the MQM position and the BSA-Seq QTL marker were 9.08 cM apart.

**Figure 4.**
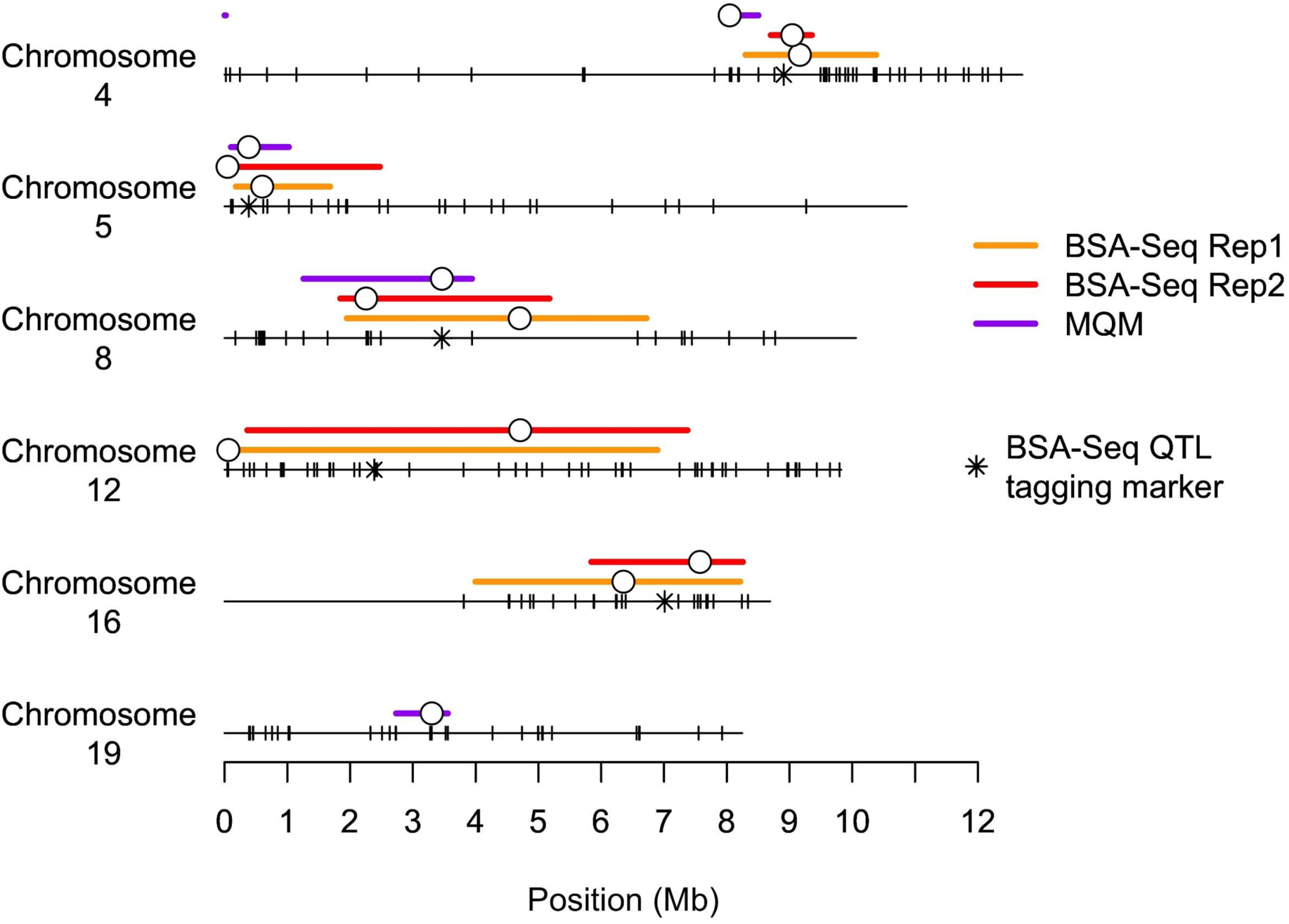
Most likely QTL positions (open circles) and credible intervals (bars) for QTL detected via BSA-Seq Reps 1 and 2 and multiple QTL mapping (MQM). Tick marks represent SNPs discovered by genotyping-by-sequencing (GBS) and used for genetic map construction. BSA-Seq QTL tagging markers were identified as those closest to the midpoint of QTL locations from Reps 1 and 2 of BSA-Seq.

**Table 4.**
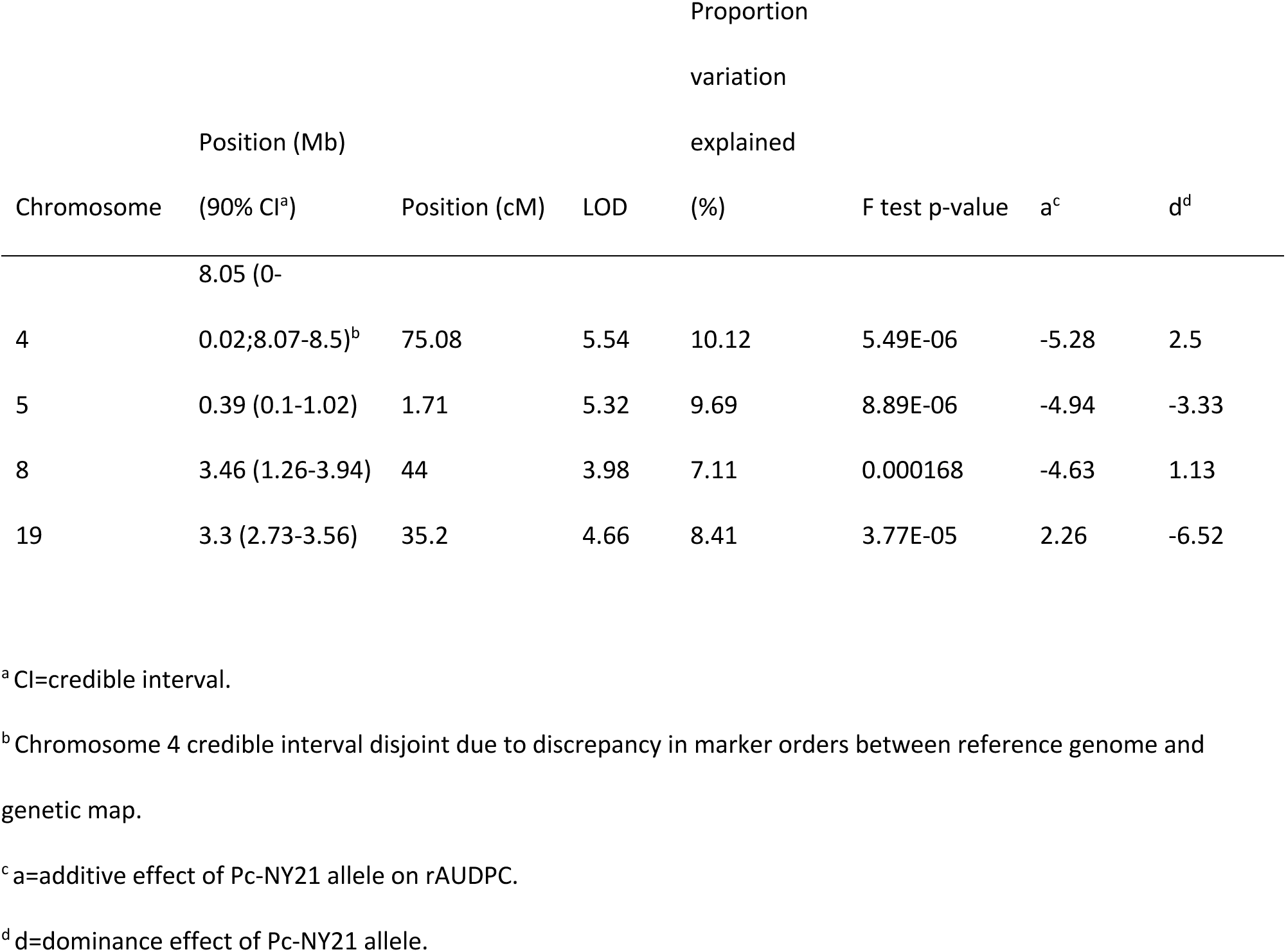
Multiple QTL mapping results and effect sizes.

### Candidate gene identification

QTL regions were defined conservatively as the union of BSA and MQM credible intervals for the purposes of candidate gene identification. Excluding the chromosome 19 QTL, which was only identified via MQM and therefore featured a much narrower credible interval, QTL regions contained between 365-815 annotated genes (Table 5). In each QTL region, between 280-548 of these genes featured variants that were polymorphic between Dunja and PcNY-21 and had a functional effect predicted to be moderate (e.g. missense mutation or in-frame insertion/deletion) or high (e.g. nonsense mutation or frameshift insertion/deletion). Of the genes with a predicted moderate or high-effect variant, 63-144 in each QTL region featured homologs in melon that were differentially expressed post inoculation with *P. capsici*. Several of these had annotations potentially related to disease resistance, including three receptor-like protein kinases on chromosomes 4, 5, and 12 (Cp4.1LG04g11730, Cp4.1LG08g07850, and Cp4.1LG12g09630, respectively) and a nucleotide-binding site Toll/interleukin-1 receptor (TIR-NB) protein on chromosome 5 (Cp4.1LG05g03340).

**Table 5.** Candidate genes in QTL regions.

| Chromosome | QTL region<br>(Mb)<sup>a</sup> | Number<br>genes | Number genes,<br>moderate effect<br>variant<sup>b</sup> | Number<br>genes, high<br>effect<br>variant<sup>b</sup> | Number<br>genes, DE<br>melon<br>homolog<sup>c</sup> | Number genes,<br>DE melon<br>homolog and<br>moderate/high<br>effect |
| --- | --- | --- | --- | --- | --- | --- |
|  | 0-0.02; |  |  |  |  |  |
| 4 | 8.07-10.38 | 365 | 234 | 46 | 99 | 63 |
| 5 | 0.04-2.47 | 425 | 205 | 33 | 127 | 63 |
| 8 | 1.25-6.72 | 666 | 444 | 104 | 175 | 115 |
| 12 | 0.06-7.38 | 815 | 415 | 65 | 234 | 127 |
| 16 | 4.00-8.26 | 694 | 404 | 85 | 217 | 144 |
| 19 | 2.73-3.56 | 85 | 32 | 5 | 23 | 11 |

<sup>a</sup> QTL region defined as union of BSA-Seq and multiple QTL mapping credible intervals.
<sup>b</sup> Functional effects of variants (moderate or high) determined by SnpEff.
<sup>c</sup> DE=differentially expressed in Wang et al. (2020); homologs between squash and melon defined as reciprocal best hits following reciprocal blastp alignments

### Prediction models

In order to assess the practical value of the QTL discovered via BSA-Seq, a multiple linear regression model containing QTL markers was compared to three genomic prediction models using genome-wide markers for their ability to predict F_2:3_ rAUDPC estimates. The median prediction accuracy of the QTL MLR model, as determined by cross-validation, was found to be moderate (0.43), although it was slightly outperformed by all three genomic prediction models. The GBLUP and Bayes-B models performed similarly (median prediction accuracies of 0.51 and 0.50, respectively). Incorporating fixed QTL marker effects into the GBLUP model, in the case of the GBLUP+QTL model, only provided a slight benefit, resulting in a median prediction accuracy of 0.54.

## Discussion

As *P. capsici* continues to spread to previously un-infested farms, often via contaminated surface water sources that may flood or inadvertently be used for irrigation (Gevens et al. 2007; Jones et al. 2014), host resistance is becoming increasingly important for the sustainable management of Phytophthora root and crown rot. Little is known, however, about the genetic basis of resistance in *C. pepo*, the squash species that includes economically important market classes such as zucchini, summer squash, and carving pumpkins. In this study, we identified a total of six genomic regions associated with variation for Phytophthora root and crown rot resistance in a biparental *C. pepo* population. Our approach, where we genotyped both individual progeny as well as bulked samples representing phenotypic extremes, also enabled us to compare the power, resolution, and practicality of two alternate strategies for discovering QTL.

Many aspects influence the QTL detection power and mapping resolution of BSA-Seq experiments. These include factors associated with the genetic architecture of the trait, such as the effect sizes, gene action, and degree of linkage between causative loci, in addition to technical considerations controlled by the experimenter, namely the size of the mapping population, size of selected bulks, and depth of sequencing coverage (Ehrenreich et al. 2010; Magwene et al. 2011; Takagi et al. 2013; Pool 2016). Generally, in a QTL mapping experiment, an increase in population size results in greater statistical power to detect QTL and more precise localization of QTL, since more recombination events are captured among the individuals in the population. However, with BSA-Seq, the gains in power and resolution conferred by an increase in population size are dependent on the choice of bulk size and sequencing coverage (Magwene et al. 2011; Pool 2016). Analytical results and simulations show that increasing the bulk proportion up to approximately 15-20% of the population size should provide an increase in power by reducing variation in allele frequencies due to random sampling (Magwene et al. 2011; Takagi et al. 2013). However, with larger population sizes, an adequate reduction in sampling variance may be achieved with lower bulk proportions (Pool 2016). Furthermore, since bulk allele frequencies are not measured directly, but are instead estimated from a sub-sample of randomly drawn sequencing reads, larger bulks may not be advantageous without corresponding increases in sequencing depth (Magwene et al. 2011; Pool 2016)

The mapping population we evaluated in this experiment, consisting of approximately 7,000 plants per rep, was large compared to most BSA-Seq experiments in plants, although several other researchers have evaluated plant populations in the range of 10,000 or even 100,000 individuals (Yang et al. 2013; Haase et al. 2015; Yuan et al. 2016). Following recommendations in the literature (Magwene et al. 2011; Takagi et al. 2013), we aimed to include 15% of the total population, or approximately 1,000 individuals, in each RAN, SUS, and RES bulk. Our realized sequencing depth of 24-70 (Table 1), however, meant that the number of reads sampled at a given site on average captured fewer than 0.5% of the approximately 2,000 distinct chromosomes represented in bulks. Consequently, variation from sequencing noise likely represented a much greater source of error in our experiment than variation from sampling individuals for bulks. Given our population size and level of sequencing depth, it may therefore have been advantageous to have selected smaller bulks, thereby applying a higher selection intensity and driving a higher divergence in allele frequency between RES and SUS bulks.

Nevertheless, we were able to overcome some of the limitations imposed by shallow sequencing and low allele frequency differentiation by using a statistical method, MULTIPOOL, that considers information from all markers on a chromosome simultaneously in order to identify the positions of QTL (Edwards and Gifford 2012). Compared to other methods for QTL mapping with BSA-Seq data, MULTIPOOL results in a low rate of false negatives and false positives even at low levels of sequencing coverage (Duitama et al. 2014; Huang et al. 2020). The two methods most popular in the plant breeding literature – QTL-Seq (Takagi et al. 2013) and the G’ test (Magwene et al. 2011), both based on sliding-window statistics – perform well in many scenarios but improperly control for multiple testing, leading to poor detection of QTL in low read depth situations (Huang et al. 2020). While the authors of MULTIPOOL do not suggest a LOD threshold for declaring a QTL significant, we were able to determine an appropriate cut-off by assessing a null distribution of LOD scores from inter-rep comparisons of bulks selected in the same direction. Our use of multiple replicates, which are not typical in BSA-Seq experiments but common in conceptually similar ‘evolve & resequence’ experiments (Long et al. 2015), also granted us greater confidence in the five QTL that were discovered, as they represented regions featuring significant differentiation between two independent selections of RES and SUS bulks.

Multiple QTL mapping with 176 F_2:3_ families resulted in the identification of a similar set of QTL compared to BSA-Seq. Both methods agreed in the identification of three regions – on chromosomes 4, 5, and 8 – where the resistant allele was inherited from resistant parent Pc-NY21. However, the two methods identified distinct loci where the allele associated with resistance was inherited from Dunja, possibly due to the lower effect sizes of these QTL. Multiple QTL mapping credible intervals were considerably narrower than the credible intervals determined by MULTIPOOL from the BSA-Seq data (Figure 3), although this is likely explained by the fact that MQM credible intervals, as estimated in R/qtl, do not account for uncertainty in the positions of other identified QTL and are therefore overly liberal (Broman and Sen 2009). Indeed, credible intervals for these same loci as identified in R/qtl via a single QTL interval mapping scan, as opposed to MQM, were similar in size to those identified via BSA-Seq (data not shown). For the loci on chromosomes 5 and 8, the most likely QTL position determined by MQM was remarkably close to the midpoint of the QTL positions identified by Reps 1 and 2 of BSA-Seq, differing by 62 kb on chromosome 5 and 16 kb on chromosome 8 (Figure 3). This was not the case, however, with the chromosome 4 QTL, where MQM determined the most likely QTL position to be over 1 Mb from the midpoint of the BSA-Seq positions. Of the 20 chromosomes, chromosome 4 also featured the greatest discrepancy in terms of marker order on the genetic and physical maps, with markers on the first 8 Mb of the chromosome inverted on the genetic map compared to their coordinates in the reference genome sequence (Figure S4). This discordance, which could either reflect a mis-assembly in the reference genome or a true inversion in the parents of our population, likely was the reason for the poor estimation of the QTL position by MULTIPOOL, as markers showing no signal in terms of allele frequency differentiation were incorrectly assumed to be tightly linked with markers featuring high signal. Several artifacts of this discrepancy between the physical and genetic maps can be seen in our results: the quite sudden allele frequency differentiation between RES and SUS pools between 8-9 Mb on chromosome 4, which appears much more abruptly compared to the onset of other QTL we identified, as well as the appearance of a second, spurious LOD peak at the beginning of the chromosome, which actually surpassed our significance threshold in Rep 2. In this case, an approach using small reference-based sliding windows, such as QTL-Seq, may have resulted in a more accurate determination of the chromosome 4 QTL position compared to a model-based method liked MULTIPOOL that considers all markers on a chromosome at once.

Overall, the two approaches – BSA-Seq using a large population of F_2_ individuals and linkage mapping with a modestly sized population of F_2:3_ families – performed similarly in terms of mapping power, resolution, and localization of QTL. Other considerations, however, may influence the decision by a researcher to choose one of these methods over the other. Genotyping of individual progeny, as required in linkage mapping, allows for the estimation of QTL effect sizes and the testing of QTL x QTL interaction effects, both of which are not possible with BSA-Seq. Individual genotyping also enables a researcher to use a single genotype dataset to map multiple traits, also impossible with BSA-Seq given that the individuals that contribute to the sequenced DNA pools are selected based on their values for one particular phenotype. On the other hand, BSA-Seq is easily scaled up to larger population sizes, making it a more powerful approach for trait mapping in early generations like an F_2_ or BC_1._ This is especially relevant in squash, since plants require manual self-pollination and take up considerable space in a field or greenhouse, making the generation of inbred lines highly resource-intensive.

Between the two QTL mapping approaches used in this experiment, a total of six QTL were identified, suggesting an oligogenic genetic architecture for Phytophthora root and crown rot resistance in squash. Each QTL was of small to moderate effect, with the largest-effect QTL we identified, those on chromosomes 4 and 5, individually accounting for only 9-10% of the phenotypic variance explained among F_2:3_ families in the MQM model. Furthermore, models including the five QTL discovered via BSA-Seq or the 4 discovered via MQM only explained a modest proportion of the phenotypic variance: 28 and 35%, respectively. Considering the relatively high broad-sense heritability for rAUDPC in the F_2:3_ population (0.77), it seems likely that we lacked the statistical power to detect additional small-effect loci contributing to variation among F_2:3_ families. Based on these results, Phytophthora root and crown resistance may have a more polygenic genetic architecture in squash compared to pepper (*Capsicum annuum*), where the genetics of resistance have been much better characterized. Crosses with different resistant pepper accessions have consistently identified, in addition to small and moderate-effect loci, a major QTL that colocalizes with a NBS-LRR gene cluster and accounts for up to 85% of the phenotypic variation explained (Thabuis et al. 2003; Thabuis et al. 2004; Quirin et al. 2005; Mallard et al. 2013; Liu et al. 2014). It remains unknown if such large-effect resistance genes exist in *Cucurbita* germplasm. In addition, mapping experiments in pepper using diverse isolates of *P. capsici* have shown that minor resistance QTL often have isolate-specific effects (Ogundiwin et al. 2004; Truong et al. 2011; Rehrig et al. 2014; Siddique et al. 2019). Although a sibling line of the resistant parent in our cross expressed a consistent resistance response when inoculated with diverse *P. capsici* isolates (LaPlant et al. 2020), it is possible that separate genes from those discovered in this experiment are involved in resistance to different isolates.

Despite the wide range of crops susceptible to Phytophthora root and crown rot (Granke et al. 2012), relatively little research has been conducted on the genetic basis of resistance in species beside pepper. Recently, however, Ramos et al. (2020) reported the identification of three resistance QTL in *C. moschata*, two of which featured resistance alleles inherited from the susceptible butternut parent of their cross. Interestingly, we found that the regions comprising the QTL identified by Ramos et al. on *C. moschata* chromosomes 11 and 14 contained blocks of shared synteny with the *C. pepo* QTL regions we identified on chromosomes 4 and 8, respectively (Figure S6). While these results are suggestive of a possible common evolutionary origin for the casual resistance genes at these loci, it also possible that QTL from the two species shared regions of synteny due to chance, especially when considering the fact that a substantial portion of the *C. pepo* genome (8%) was contained within the six QTL we identified. Because Ramos et al. used a BSA-Seq strategy combined with further selective genotyping for marker validation, QTL effect sizes were not reported, making additional comparisons between QTL from the two experiments difficult.

Unfortunately, the successful identification of candidate genes in our experiment was made difficult by the high degree of statistical uncertainty in QTL positions, as the intersections of credible intervals from BSA-Seq and MQM generally spanned several Mb and comprised hundreds of genes (Figure 4; Table 5). Furthermore, unlike qualitative disease resistance, which is typically conferred by proteins belonging to a small number of well-characterized gene families (Kourelis and van der Hoom 2018), quantitative disease resistance in plants is mediated by a wide variety of genes with diverse functions (Nelson et al. 2018). In squash, Phytophthora root and crown rot resistance is associated with a reduction in pathogen infection of vascular tissue (Krasnow et al. 2017), similar to in pepper, where the secretion of root exudates and the formation of callose and cell well appositions appear to prevent the colonization of *P. capsici* beyond the outermost layers of the cortex in resistant roots and stems (Kim and Kim 2009; Dunn and Smart 2015; Piccini et al. 2019). The genes that confer these defense responses are unknown and may have various functions, such as those related to pathogen recognition, signaling, or the production of antimicrobial molecules. Therefore, we found functional annotations relatively uninformative for the identification of candidate genes in QTL intervals. However, we were able to identify a reduced number of higher-confidence candidate genes – between 63-144 per QTL region – that were segregating for variants of moderate or high effect in our population and that featured a homolog in melon that was differentially expressed post inoculation with *P. capsici*. Additional sources of information, such as transcriptomic data on the parents of our population, may be able to further prioritize candidate genes in QTL intervals.

The results from this experiment are promising for the implementation of MAS for Phytophthora root and crown rot resistance in squash. Although QTL effect sizes were small, we showed that prediction models using just five markers – one for each QTL detected via BSA-seq – could predict the resistance levels of F_2:3_ families with a drop in prediction accuracy of only 0.07-0.08 compared to whole genome prediction models that did not fit QTL effects (Figure 3). While this difference is not negligible, genotyping progeny at five markers instead of several hundred may be more easy to implement in a breeding program and could translate to meaningful cost savings. Nevertheless, further research is still necessary to test the effects of these QTL in different populations and environments. It should be noted that both parents of our cross belonged to *C. pepo* ssp. *pepo*, the more resistant (Camp et al. 2009; Meyer and Hausbeck 2012; Krasnow et al. 2017) of the two independently domesticated subspecies of *C. pepo* (Decker 1988; Sanjur et al. 2002) and were therefore both likely fixed for the resistant alleles at additional, unknown loci that are presumably polymorphic between *C. pepo* ssp. *pepo* and ssp. *ovifera*. Efforts to introgress resistance from Pc-NY21 into *C. pepo* ssp. *ovifera* cultivars via MAS may not be successful without accounting for these loci. Furthermore, it is important to mention that neither of the parents of our mapping population were fully inbred lines, since they were chosen more for their utility in breeding rather than their suitability for QTL mapping. By only using informative markers that were homozygous in the parents, we were able to map the genomic positions of QTL that were polymorphic between Dunja and Pc-NY21. If the parent carrying the resistant allele for a given QTL was actually heterozygous at that locus, however, these markers would be uninformative for MAS, as they would be unable to differentiate between the resistant and susceptible haplotypes from the donor parent of the resistant allele. We suspect, however, that this is not the case for the major QTL we identified on chromosomes 4, 5, and 8, given the low proportion of heterozygous markers in PcNY-21 in these regions (data not shown).

In conclusion, we discovered a total of six genomic regions associated with Phytophthora root and crown rot resistance in a biparental squash population. Three featured resistant alleles inherited from the resistant parent of our cross and were identified independently via both BSA-Seq and linkage mapping; the other three featured resistant alleles inherited from the susceptible parent of the cross and were each identified via only one of the two methods. Despite the small to moderate effect sizes of these QTL, we believe that markers in these regions can be used to accelerate disease resistance breeding in various market classes of squash. Our mapping resolution was too low to conclusively identify strong candidates for genes conferring resistance in QTL regions. Nevertheless, these results represent an early step toward uncovering the genetic and molecular mechanisms underlying Phytophthora root and crown rot resistance in squash.

## Acknowledgements

We are grateful for technical assistance from Andrew Aldcroft, Mariami Bekauri, Colin Day, Garrett Giles, Holly Lange, Nicholas King, Carolina Puentes Silva, and Carolina Vogel, as well as statistical advice from Chris Hernandez and Ryokei Tanaka. We would also like to thank Jeff Glaubitz, Peter Schweitzer, and Jing Wu from the Cornell Institute of Biotechnology for consultation regarding library preparation for whole genome sequencing; Mainor Najera, Fernando Villalta Mata, and Andrés Villalta Pereira from VillaPlants for provision of seed increase services; and Mary Kreitinger for assistance in obtaining permits for shipping of seed, tissue, and supplies. Funding for this project was provided by the Cucurbit Coordinated Agricultural Project (CucCAP; USDA National Institute of Food and Agriculture Specialty Crop Research Initiative Grant No. 2015-51181-24285) and the New York State Department of Agriculture and Markets.

---

**Figure S1.**
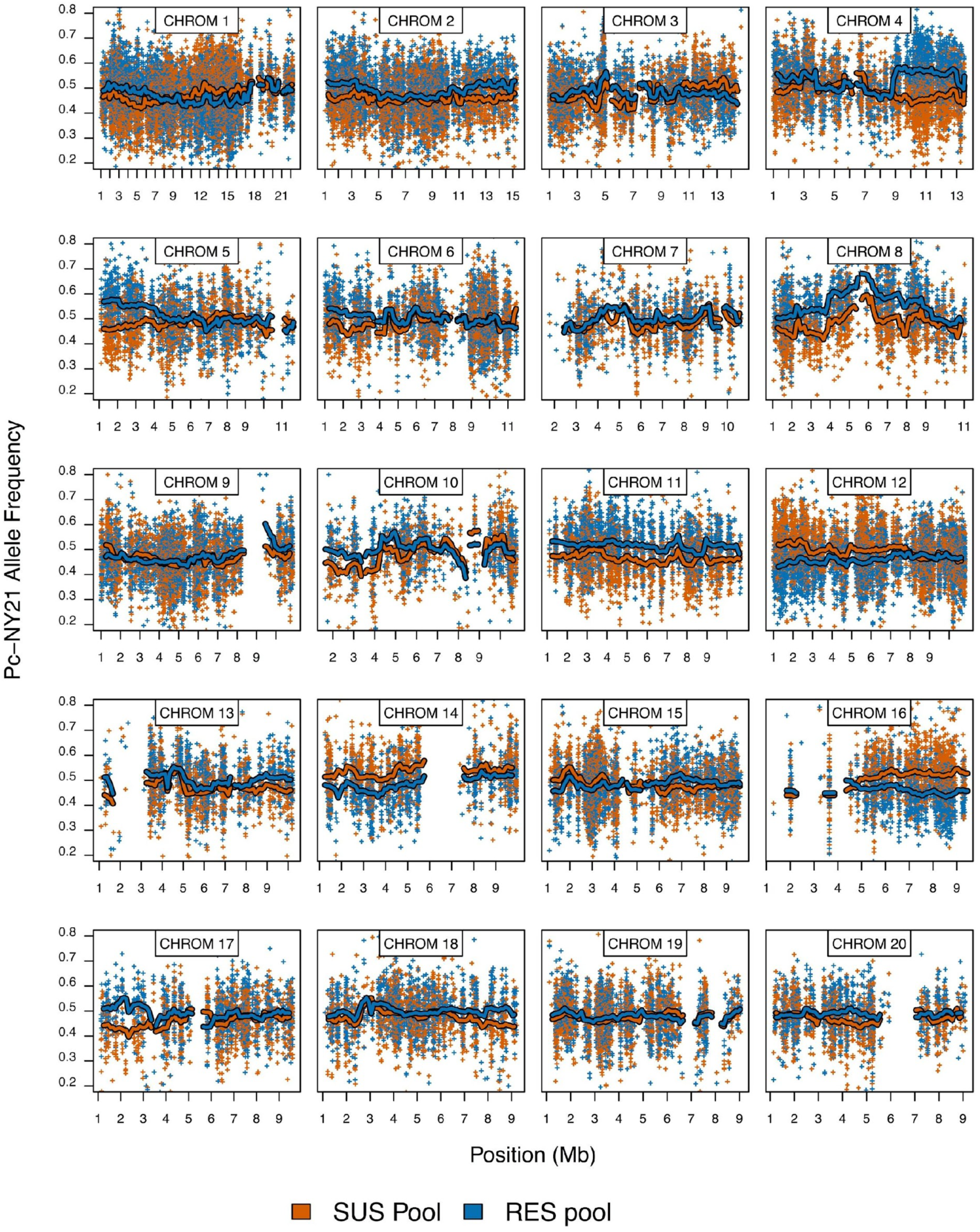
Pc-NY21 allele frequencies in Rep 1 susceptible (SUS) and resistant (RES) pools across all 20 chromosomes. Plus signs are individual SNP allele frequencies and lines are smoothed means calculated in 500 Kb sliding windows with a 100 Kb increment. For simplicity of visualization, only a random subset of 25% of SNPs are shown. Smoothed means are not calculated in any window featuring fewer than 30 SNPs.

**Figure S2.**
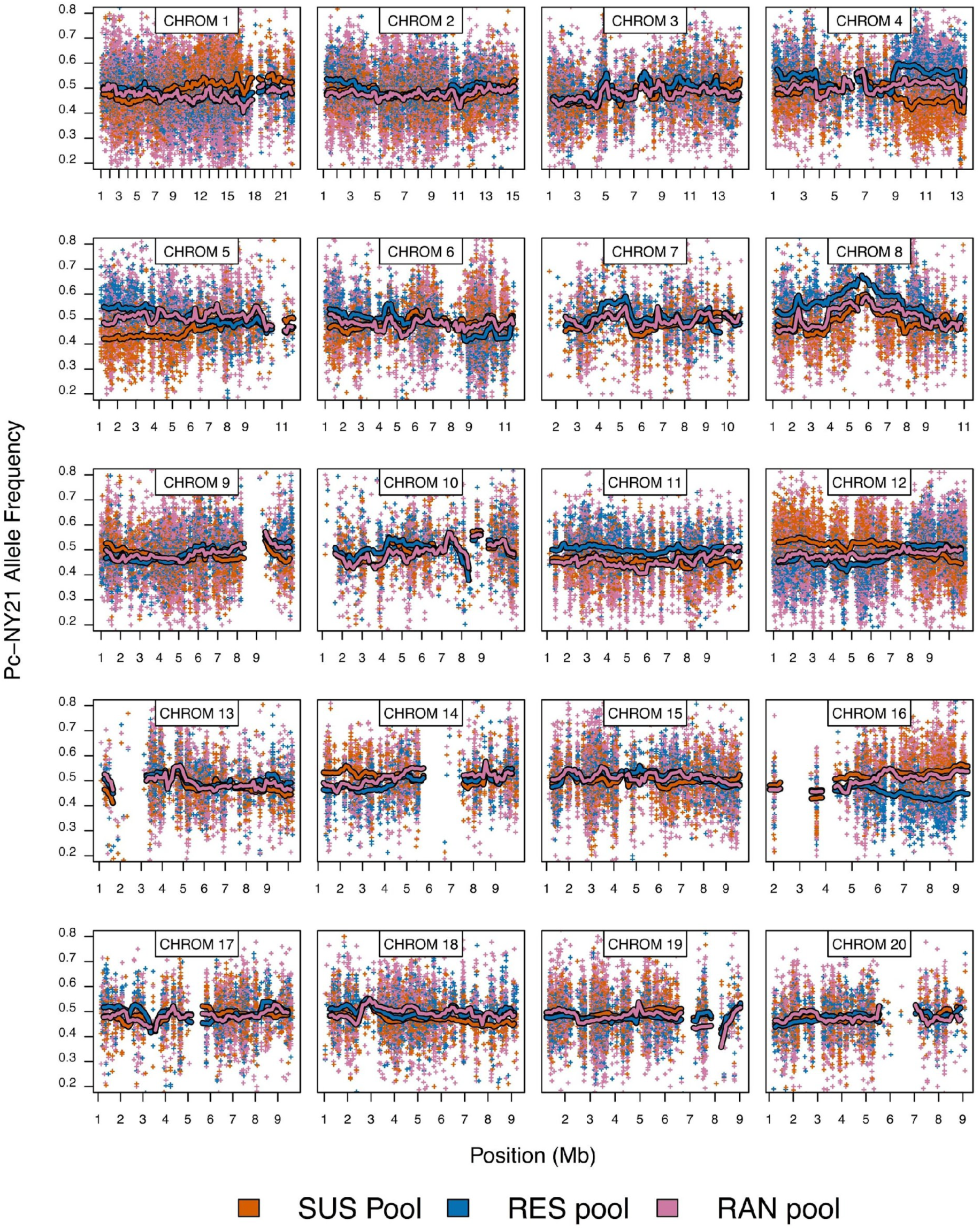
Pc-NY21 allele frequencies in Rep 2 susceptible (SUS), resistant (RES), and random (RAN) pools across all 20 chromosomes. Plus signs are individual SNP allele frequencies and lines are smoothed means calculated in 500 Kb sliding windows with a 100 Kb increment. For simplicity of visualization, only a random subset of 25% of SNPs are shown. Smoothed means are not calculated in any window featuring fewer than 30 SNPs.

**Figure S3.**
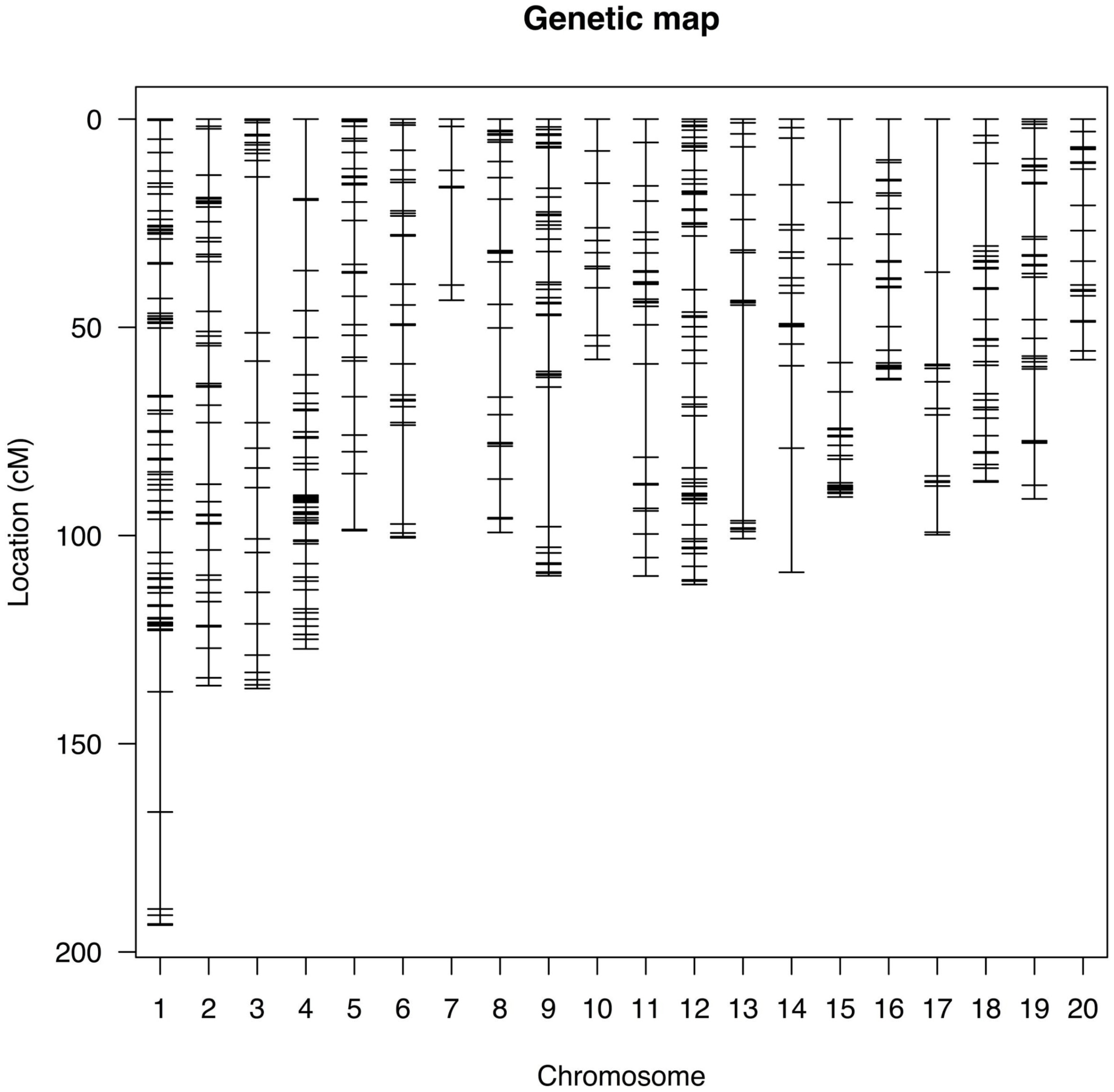
Position of 605 SNP markers on 20 linkage groups of a genetic map created using 181 F_2_ individuals.

**Figure S4.**
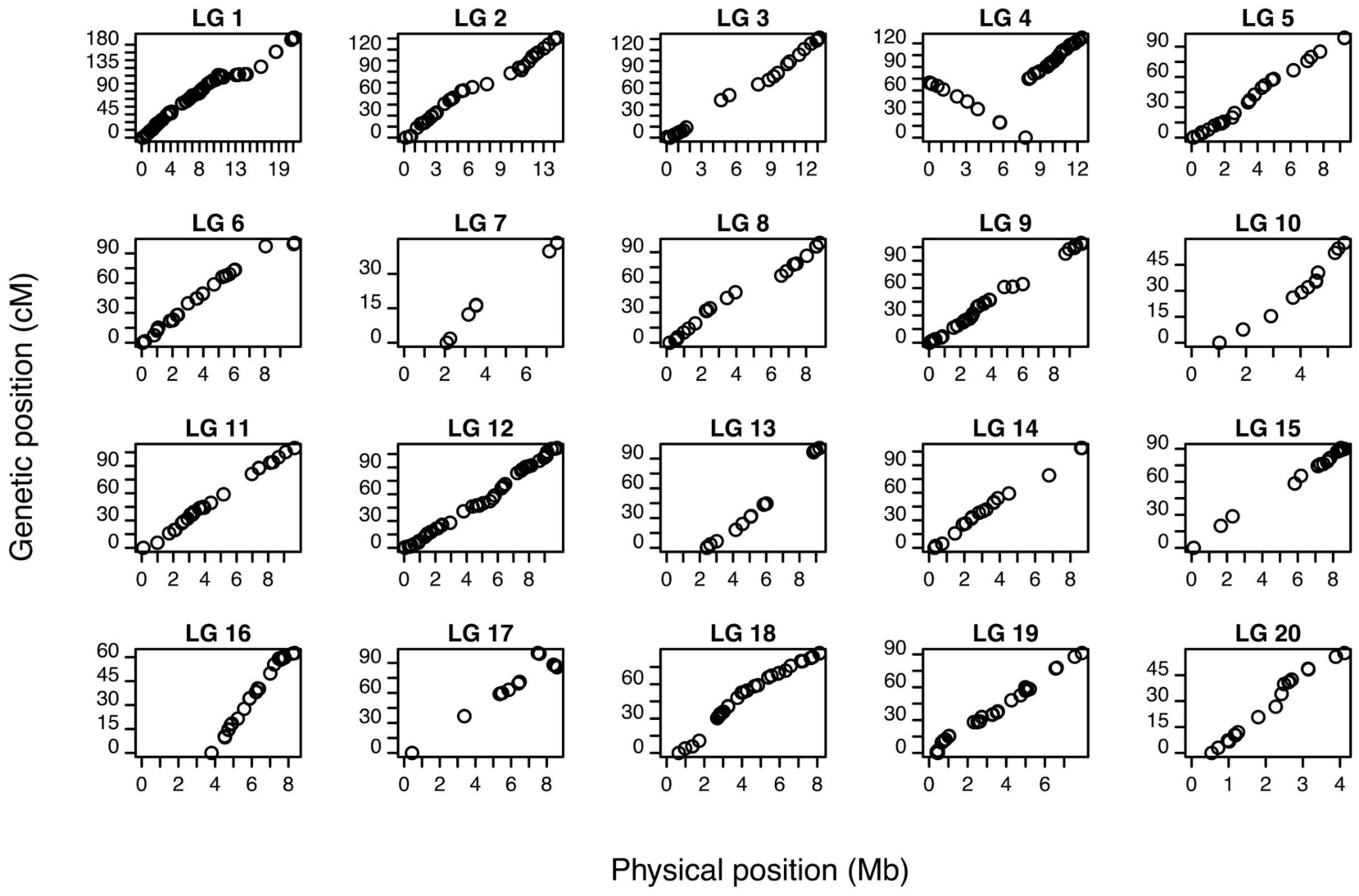
Physical position (Mb) vs genetic position (cM) of 605 SNP markers placed on the genetic map. Markers assigned to a different linkage group on the genetic map compared to their location in the reference genome are not shown.

**Figure S5.**
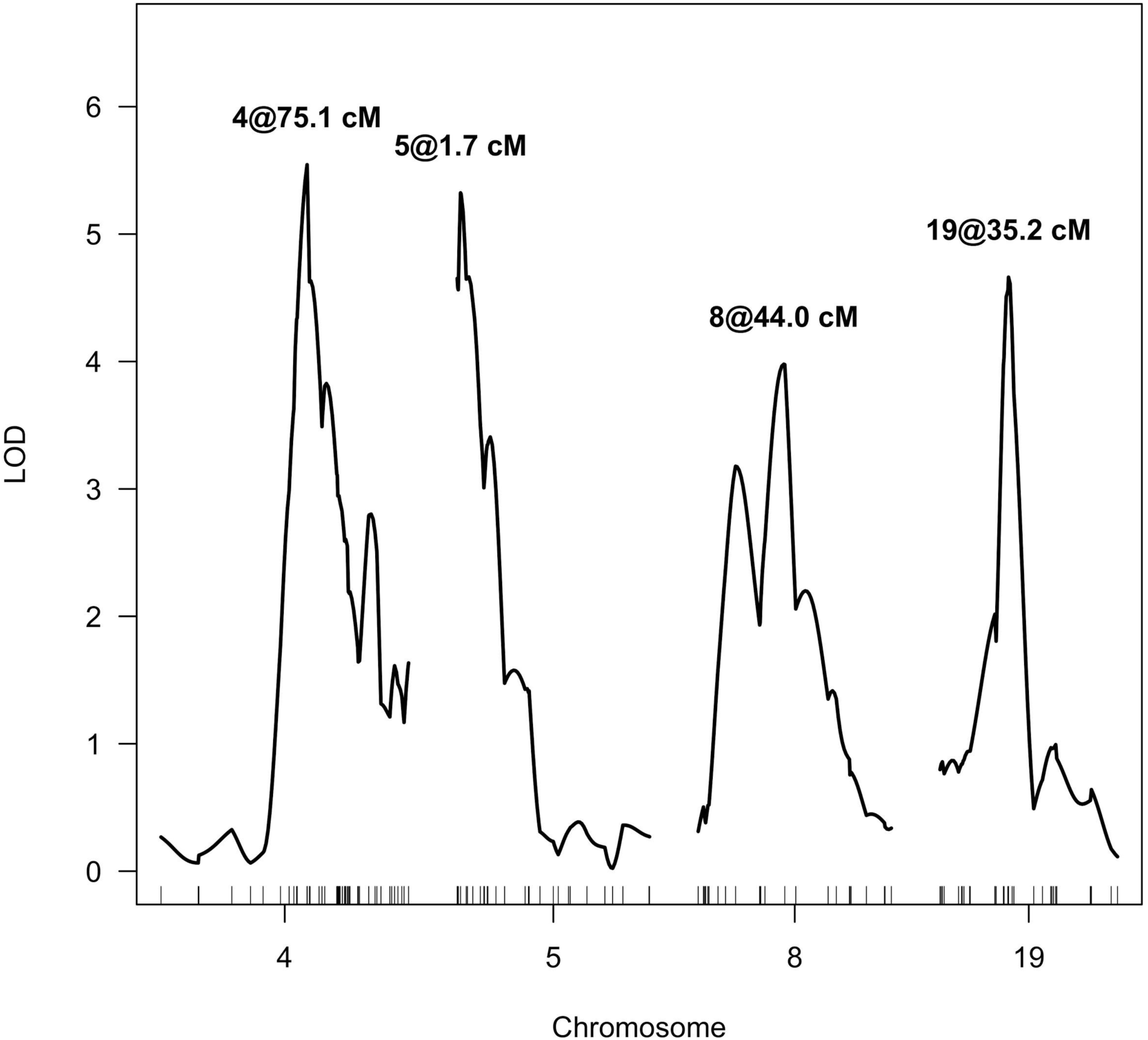
Multiple QTL mapping LOD scores using rAUDPC estimates for 176 F_2:3_ families and a genetic map with 605 SNPs called on their F_2_ parents. Only linkage groups where a QTL was identified are shown.

**Figure S6.**
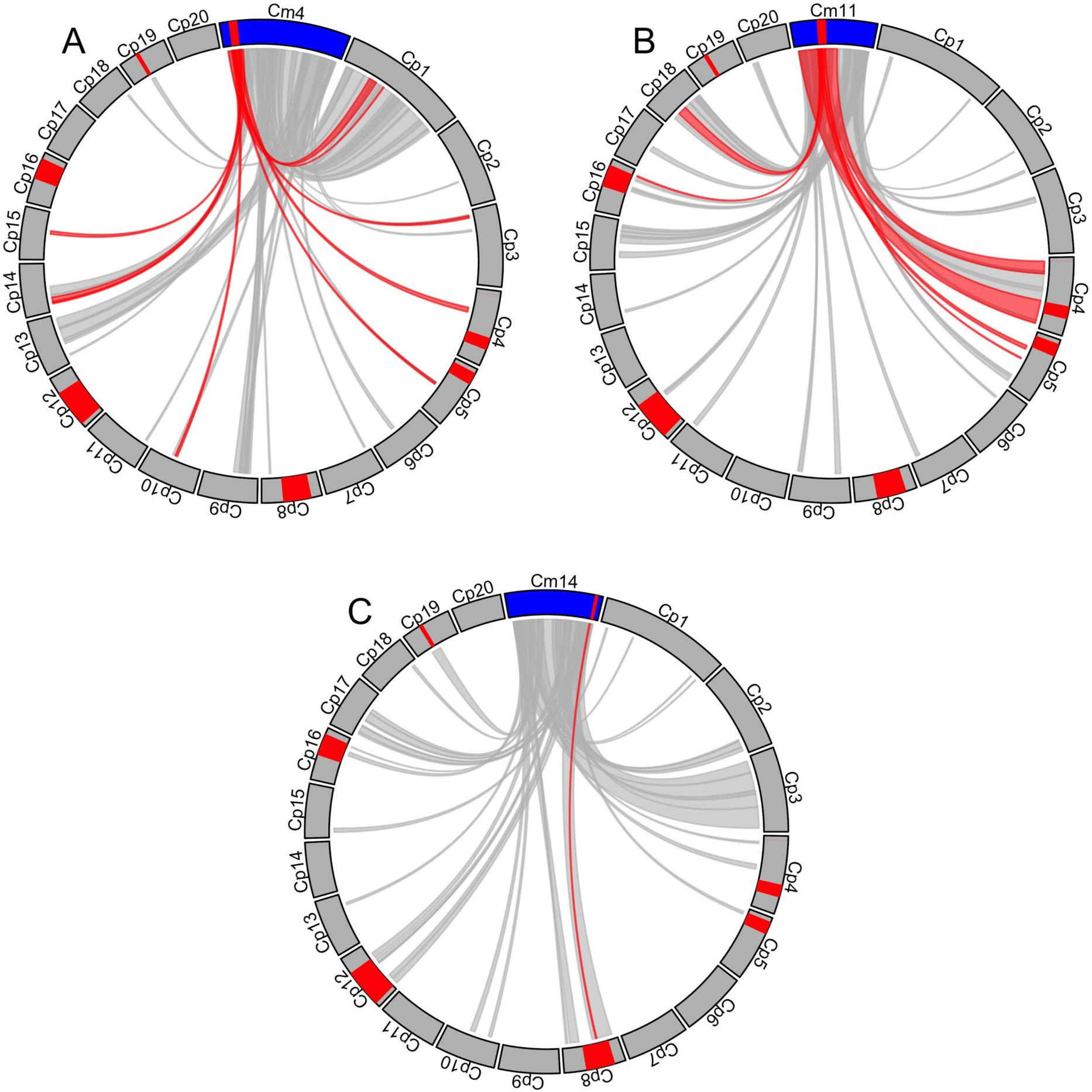
Syntenic blocks shared between *C. moschata* chromosomes 4, 11, and 14 (A-C, respectively) and *C. pepo*. QTL regions identified in *C. moschata* by Ramos et al. (2020) and in *C. pepo* in this publication are colored in red. Chords between syntenic blocks where the *C. moschata* counterpart overlaps a QTL are colored in red. Only syntenic blocks with a score >400 as reported by MCScanX are shown.

**Table S1.** Broad-sense heritabilities (*H^2^*), parental and F_1_ estimates, and medians of ‘random’ and ‘selected’ cohorts of F_2:3_ families for five traits.

| Trait<sup>a</sup> | $H^2$ | Dunja | PcNY-21 | Dunja x | Median,<br>'random' F<sub>2:3</sub> | Median,<br>'selected' F<sub>2:3</sub> |
| --- | --- | --- | --- | --- | --- | --- |
|  |  |  |  | PcNY-21 F<sub>1</sub> | families | families |
| dpi3 | 0.4 | 16.67 | 0 | 25 | 19.44 | 12.31 |
| dpi5 | 0.66 | 74.07 | 13.89 | 54.17 | 63.64 | 45.04 |
| dpi7 | 0.72 | 94.44 | 41.67 | 75 | 83.33 | 61.25 |
| dpi10 | 0.76 | 100 | 55.56 | 91.67 | 97.22 | 86.11 |
| rAUDPC | 0.77 | 80.95 | 32.61 | 68.54 | 74.18 | 60.59 |

<sup>a</sup> dpi3, dpi5, dpi7, dpi10 = plot mortality at 3, 5, 7, and 10 days post inoculation, respectively. rAUDPC = relative Area Under the Disease Progress Curve.

Table S2. Genetic map positions of 605 SNPs called on 181 F_2_ individuals.

**Table S3.** Positions of SNP markers tagging BSA-Seq QTL and their PcNY-21 allele frequencies in ‘random’ and ‘selected’ F_2_ cohorts.

| Chromosome | Position (bp) | Distance from BSA<br>mean LOD peak<br>(bp) | PcNY-21 allele<br>frequency,<br>'random' $F_{2:3}$<br>families | PcNY-21 allele<br>frequency,<br>'selected' $F_{2:3}$<br>families |
| --- | --- | --- | --- | --- |
| 4 | 8908173 | 195977 | 0.47 | 0.66 |
| 5 | 385710 | 62160 | 0.51 | 0.63 |
| 8 | 3462310 | 15990 | 0.53 | 0.71 |
| 12 | 2386018 | 968 | 0.54 | 0.42 |
| 16 | 7011538 | 47288 | 0.46 | 0.42 |

Table S4. Annotations for all genes annotated in QTL regions, as well as their number of predicted moderate or high effect variants and whether they feature a differentially expressed (DE) homolog in melon (NA if no homolog identified).

## References

1. Bates D, Mächler M, Bolker B, Walker S (2015) Fitting linear mixed-effects models using lme4. J Stat Softw 67:1–48

2. Bolger AM, Lohse M, Usadel B (2014) Trimmomatic: a flexible trimmer for Illumina sequence data. Bioinformatics 30:2114–2120

3. Broman KW, Sen Ś (2009) Fit and exploration of multiple-QTL models. In: Broman KW, Sen S (eds) A Guide to QTL Mapping with R/qtl. Springer, New York, NY, pp 241–282

4. Broman KW, Wu H, Sen Ś, Churchill GA (2003) R/qtl: QTL mapping in experimental crosses. Bioinformatics 19:889–890

5. Camacho C, Coulouris G, Avagyan V, Ma N, Papadopoulos J, Bealer K, Madden TL (2009) BLAST+: architecture and applications. BMC Bioinformatics 10:421

6. Camp AR, Lange HW, Reiners S, Dillard HR, Smart CD (2009) Tolerance of summer and winter squash lines to Phytophthora blight, 2008. Plant Disease Management Reports 3:V022

7. Chavez DJ, Kabelka EA, Chaparro JX (2011) Screening of *Cucurbita moschata* Duchesne germplasm for crown rot resistance to Floridian isolates of *Phytophthora capsici* Leonian. HortScience 46:536–540

8. Chen S, Zhou Y, Chen Y, Gu J (2018) fastp: an ultra-fast all-in-one FASTQ preprocessor. Bioinformatics 34:i884–i890

9. Cingolani P, Platts A, Wang LL, Coon M, Nguyen T, Wang L, Land SJ, Lu X, Ruden DM (2012) A program for annotating and predicting the effects of single nucleotide polymorphisms, SnpEff. Fly 6:80–92

10. Danecek P, Auton A, Abecasis G, Albers CA, Banks E, DePristo MA, Handsaker RE, Lunter G, Marth GT, Sherry ST, McVean G, Durbin R (2011) The variant call format and VCFtools. Bioinformatics 27:2156–2158

11. Decker DS (1988) Origin(s), evolution, and systematics of *Cucurbita pepo* (Cucurbitaceae). Econ Bot 42:4–15

12. Dunn AR, Milgroom MG, Meitz JC, McLeod A, Fry WE, McGrath MT, Dillard HR, Smart CD (2010) Population structure and resistance to mefenoxam of *Phytophthora capsici* in New York State. Plant Dis 94:1461–1468

13. Dunn AR, Smart CD (2015) Interactions of *Phytophthora capsici* with resistant and susceptible pepper roots and stems. Phytopathology 105:1355–1361

14. Edwards MD, Gifford DK (2012) High-resolution genetic mapping with pooled sequencing. BMC Bioinformatics 13:S8

15. Elshire RJ, Glaubitz JC, Sun Q, Poland JA, Kawamoto K, Buckler ES, Mitchell SE (2011) A robust, simple genotyping-by-sequencing (GBS) approach for high diversity species. PLoS ONE 6:e19379

16. Endelman JB (2011) Ridge regression and other kernels for genomic selection with R Package rrBLUP. Plant Genome 4:250–255

17. Enzenbacher TB, Hausbeck MK (2012) An evaluation of cucurbits for susceptibility to Cucurbitaceous and Solanaceous *Phytophthora capsici* isolates. Plant Dis 96:1404–1414

18. Esteras C, Gómez P, Monforte AJ, Blanca J, Vicente-Dólera N, Roig C, Nuez F, Picó B (2012) High-throughput SNP genotyping in *Cucurbita pepo* for map construction and quantitative trait loci mapping. BMC Genomics 13:80

19. Fry WE (1978) Quantification of general resistance of potato cultivars and fungicide effects for integrated control of potato late blight. Phytopathology 68:1650

20. Garcia-Mas J, Benjak A, Sanseverino W, et al (2012) The genome of melon (*Cucumis melo* L.). Proc Natl Acad Sci USA 109:11872–11877

21. Gevens AJ, Donahoo RS, Lamour KH, Hausbeck MK (2007) Characterization of *Phytophthora capsici* from Michigan surface irrigation water. Phytopathology 97:421–428.

22. Granke LL, Quesada-Ocampo L, Lamour K, Hausbeck MK (2012) Advances in research on *Phytophthora capsici* on vegetable crops in the United States. Plant Dis 96:1588–1600

23. Gu Z, Gu L, Eils R, Schlesner M, Brors B (2014) *circlize* implements and enhances circular visualization in R. Bioinformatics 30:2811–2812

24. Haley CS, Knott SA (1992) A simple regression method for mapping quantitative trait loci in line crosses using flanking markers. Heredity 69:315–324

25. Hausbeck MK, Lamour KH (2004) *Phytophthora capsici* on vegetable crops: research progress and management challenges. Plant Dis 88:1292–1303

26. Holdsworth WL, LaPlant KE, Bell DC, Jahn MM, Mazourek M (2016) Cultivar-based introgression mapping reveals wild species-derived Pm-0, the major powdery mildew resistance locus in squash. PLoS ONE 11:e0167715

27. Holland JB, Nyquist WE, Cervantes-Martínez CT (2003) Estimating and interpreting heritability for plant breeding: an update. In: Plant Breeding Reviews. John Wiley & Sons, Ltd, pp 9–112

28. Huang L, Tang W, Bu S, Wu W (2020) BRM: a statistical method for QTL mapping based on bulked segregant analysis by deep sequencing. Bioinformatics 36:2150–2156

29. Illa-Berenguer E, Van Houten J, Huang Z, van der Knaap E (2015) Rapid and reliable identification of tomato fruit weight and locule number loci by QTL-seq. Theor Appl Genet 128:1329–1342

30. Jackson KL, Yin J, Ji P (2012) Sensitivity of *Phytophthora capsici* on vegetable crops in Georgia to mandipropamid, dimethomorph, and cyazofamid. Plant Dis 96:1337–1342

31. Jones LA, Worobo RW, Smart CD (2014) Plant-pathogenic oomycetes, *Escherichia coli* strains, and *Salmonella* spp. frequently found in surface water used for irrigation of fruit and vegetable crops in New York State. Appl Environ Microbiol 80:4814–4820

32. Kim S-G, Kim Y-H (2009) Histological and cytological changes associated with susceptible and resistant responses of chili pepper root and stem to *Phytophthora capsici* infection. Plant Pathol J 25:113–120

33. Kourelis J, van der Hoorn RAL (2018) Defended to the nines: 25 years of resistance gene cloning identifies nine mechanisms for R protein function. Plant Cell 30:285–299

34. LaPlant KE, Vogel G, Reeves E, Smart CD, Mazourek M (2020) Performance and resistance to phytophthora crown and root rot in squash lines. HortTechnology 1:1–11 https://doi.org/10.21273/HORTTECH04636-20

35. Lamour KH, Hausbeck MK (2000) Mefenoxam insensitivity and the sexual stage of *Phytophthora capsici* in Michigan cucurbit fields. Phytopathology 90:396–400

36. Lee BK, Kim BS, Chang SW, Hwang BK (2001) Aggressiveness to pumpkin cultivars of isolates of *Phytophthora capsici* from pumpkin and pepper. Plant Dis 85:497–500

37. Li H, Durbin R (2009) Fast and accurate short read alignment with Burrows–Wheeler transform. Bioinformatics 25:1754–1760

38. Li H, Handsaker B, Wysoker A, et al (2009) The Sequence Alignment/Map format and SAMtools. Bioinformatics 25:2078–2079

39. Liu W-Y, Kang J-H, Jeong H-S, Choi H-J, Yang H-B, Kim K-T, Choi D, Choi GJ, Jahn M, Kang B-C (2014) Combined use of bulked segregant analysis and microarrays reveals SNP markers pinpointing a major QTL for resistance to *Phytophthora capsici* in pepper. Theor Appl Genet 127:2503–2513

40. Long A, Liti G, Luptak A, Tenaillon O (2015) Elucidating the molecular architecture of adaptation via evolve and resequence experiments. Nat Rev Genet 16:567–582

41. Lust TA, Paris HS (2016) Italian horticultural and culinary records of summer squash (*Cucurbita pepo*, Cucurbitaceae) and emergence of the zucchini in 19th-century Milan. Ann Bot 118:53–69

42. Magwene PM, Willis JH, Kelly JK (2011) The statistics of bulk segregant analysis using next generation sequencing. PLoS Comput Biol 7:e1002255

43. Mallard S, Cantet M, Massire A, Bachellez A, Ewert S, Lefebvre V (2013) A key QTL cluster is conserved among accessions and exhibits broad-spectrum resistance to *Phytophthora capsici*: a valuable locus for pepper breeding. Mol Breed 32:349–364

44. Melo ATO, Bartaula R, Hale I (2016) GBS-SNP-CROP: a reference-optional pipeline for SNP discovery and plant germplasm characterization using variable length, paired-end genotyping-by-sequencing data. BMC Bioinformatics 17:29

45. Mendiburu FD, Simon R (2015) Agricolae -Ten years of an open source statistical tool for experiments in breeding, agriculture and biology. PeerJ PrePrints

46. Menezes CB, Maluf WR, Faria MV, Azevedo SM, Resende JT, Figueira AR, Gomes LA (2015) Inheritance of resistance to papaya ringspot virus-watermelon strain (PRSV-W) in “Whitaker” summer squash line. Crop Breeding and Applied Biotechnology 15:203–209

47. Meuwissen THE, Hayes BJ, Goddard ME (2001) Prediction of total genetic value using genome-wide dense marker maps. Genetics 157:1819–1829

48. Michael VN, Fu Y, Meru G (2019) Inheritance of resistance to phytophthora crown rot in *Cucurbita pepo*. HortScience 54:1156–1158

49. Michelmore RW, Paran I, Kesseli RV (1991) Identification of markers linked to disease-resistance genes by bulked segregant analysis: a rapid method to detect markers in specific genomic regions by using segregating populations. Proc Natl Acad Sci USA 88:9828–9832

50. Montero-Pau J, Blanca J, Bombarely A, Ziarsolo P, Esteras C, Martí-Gomez C, Ferriol M, Gómez P, Jamilena M, Mueller L, Picó B, Cañizares J (2018) De novo assembly of the zucchini genome reveals a whole-genome duplication associated with the origin of the *Cucurbita* genus. Plant Biotechnol J 16:1161–1171

51. Montero-Pau J, Blanca J, Esteras C, Martínez-Pérez EM, Gómez P, Monforte AJ, Cañizares J, Picó B. (2017) An SNP-based saturated genetic map and QTL analysis of fruit-related traits in zucchini using genotyping-by-sequencing. BMC Genomics 18:94

52. Nelson R, Wiesner-Hanks T, Wisser R, Balint-Kurti P (2018) Navigating complexity to breed disease-resistant crops. Nat Rev Genet 19:21–33

53. Ogundiwin EA, Berke TF, Massoudi M, Black LL, Huestis G, Choi D, Lee S, Prince JP (2005) Construction of 2 intraspecific linkage maps and identification of resistance QTLs for *Phytophthora capsici* root-rot and foliar-blight diseases of pepper (*Capsicum annuum* L.). Genome 48:698–711

54. Padley LD, Kabelka EA, Roberts PD (2009) Inheritance of resistance to crown rot caused by *Phytophthora capsici* in *Cucurbita*. HortScience 44:211–213

55. Padley LD, Kabelka EA, Roberts PD, French R (2008) Evaluation of *Cucurbita pepo* accessions for crown rot resistance to isolates of *Phytophthora capsici*. HortScience 43:1996–1999

56. Parra G, Ristaino JB (2001) Resistance to mefenoxam and metalaxyl among field isolates of *Phytophthora capsici* causing Phytophthora blight of bell pepper. Plant Dis 85:1069–1075

57. Pérez P, Campos G de los (2014) Genome-wide regression and prediction with the BGLR statistical package. Genetics 198:483–495

58. Piccini C, Parrotta L, Faleri C, Romi M, Del Duca S, Cai G (2019) Histomolecular responses in susceptible and resistant phenotypes of *Capsicum annuum* L. infected with *Phytophthora capsici*. Sci Hortic 244:122–133

59. Pool JE (2016) Genetic mapping by bulk segregant analysis in *Drosophila*: experimental design and simulation-based inference. Genetics 204:1295–1306

60. Quinlan AR, Hall IM (2010) BEDTools: a flexible suite of utilities for comparing genomic features. Bioinformatics 26:841–842

61. Quirin EA, Ogundiwin EA, Prince JP, Mazourek M, Briggs MO, Chlanda TS, Kim KT, Falise M, Kang BC, Jahn MM (2005) Development of sequence characterized amplified region (SCAR) primers for the detection of Phyto.5.2, a major QTL for resistance to *Phytophthora capsici* Leon. in pepper. Theor Appl Genet 110:605–612

62. R Core Team (2019) R: A langauge and environment for statistical computing. Vienna, Austria

63. Ramos A, Fu Y, Michael V, Meru G (2020) QTL-seq for identification of loci associated with resistance to Phytophthora crown rot in squash. Sci Rep 10:5326

64. Rehrig WZ, Ashrafi H, Hill T, Prince J, Van Deynze A (2014) CaDMR1 cosegregates with QTL Pc5.1 for resistance to *Phytophthora capsici* in pepper (*Capsicum annuum*). Plant Genome 7:1–12

65. Rhodes AM (1964) Inheritance of powdery mildew resistance in the genus *Cucurbita*. Plant Dis Rep 48:54–55

66. Sanjur OI, Piperno DR, Andres TC, Wessel-Beaver L (2002) Phylogenetic relationships among domesticated and wild species of *Cucurbita* (Cucurbitaceae) inferred from a mitochondrial gene: implications for crop plant evolution and areas of origin. Proc Natl Acad Sci USA 99:535– 540

67. Siddique MI, Lee H-Y, Ro N-Y, Han K, Venkatesh J, Solomon AM, Patil AS, Changkwian AC, Kwon J-K, Kang B-C (2019) Identifying candidate genes for *Phytophthora capsici* resistance in pepper (*Capsicum annuum*) via genotyping-by-sequencing-based QTL mapping and genome-wide association study. Sci Rep 9:9962

68. Sun H, Wu S, Zhang G, et al (2017) Karyotype stability and unbiased fractionation in the paleo-allotetraploid *Cucurbita* genomes. Mol Plant 10:1293–1306.

69. Takagi H, Abe A, Yoshida K, et al (2013) QTL-seq: rapid mapping of quantitative trait loci in rice by whole genome resequencing of DNA from two bulked populations. Plant J 74:174–183

70. Taylor J, Butler D (2017) R package ASMap: efficient genetic linkage map construction and diagnosis. J Stat Soft 79:1–29

71. Thabuis A, Palloix A, Pflieger S, Daubéze A-M, Caranta C, Lefebvre V (2003) Comparative mapping of *Phytophthora* resistance loci in pepper germplasm: evidence for conserved resistance loci across Solanaceae and for a large genetic diversity. Theor Appl Genet 106:1473–1485

72. Thabuis A, Palloix A, Servin B, Daubéze AM, Signoret P, Hospital F, Lefebvre V (2004) Marker-assisted introgression of 4 *Phytophthora capsici* resistance QTL alleles into a bell pepper line: validation of additive and epistatic effects. Mol Breed 14:9–20

73. Tian D, Babadoost M (2004) Host range of *Phytophthora capsici* from pumpkin and pathogenicity of isolates. Plant Dis 88:485–489

74. Truong HTH, Kim KT, Kim DW, Kim S, Chae Y, Park JH, Oh DG, Cho MC (2012) Identification of isolate-specific resistance QTLs to phytophthora root rot using an intraspecific recombinant inbred line population of pepper (*Capsicum annuum*). Plant Pathol 61:48–56

75. United States Department of Agriculture (2019) Vegetables and melons. In: Agricultural Statistics 2019. United States Government Printing Office, Washington, D.C.

76. Wang Y, Tang H, Debarry JD, et al (2012) MCScanX: a toolkit for detection and evolutionary analysis of gene synteny and collinearity. Nucleic Acids Res 40:e49

77. Wang P, Wu H, Zhao G, He Y, Kong W, Zhang J, Liu S, Liu M, Hu K, Liu L, Xu Y, Xu Z (2020) Transcriptome analysis clarified genes involved in resistance to *Phytophthora capsici* in melon. PLoS ONE 15:e0227284

78. Wu Y, Bhat PR, Close TJ, Lonardi S (2008) Efficient and accurate construction of genetic linkage maps from the minimum spanning tree of a graph. PLoS Genet 4:e1000212

79. Zheng Y, Wu S, Bai Y, et al (2019) Cucurbit Genomics Database (CuGenDB): a central portal for comparative and functional genomics of cucurbit crops. Nucleic Acids Res 47:D1128–D1136

